# Representation of *Drosophila* larval behaviors by muscle activity patterns

**DOI:** 10.1101/2021.11.26.470133

**Authors:** Jinrun Zhou, Zenan Huang, Xinhang Li, Zhiying Song, Yixuan Sun, Junyu Ping, Xiaopeng Chen, Peng Fei, Nenggan Zheng, Zhefeng Gong

## Abstract

How muscle actions are coordinated to realize animal movement is a fundamental question in behavioral study. To obtain the overall muscular activity patterns accompanying behaviors at high spatiotemporal resolution is technically difficult. In this work, we used light sheet microscopy to simultaneously image and analyze the activity, length and orientation of *Drosophila* larval muscles across body segments at single muscle resolution in nearly free behaviors. For typical behavioral modes such as peristalsis, head cast and turning, larval muscles showed behavioral mode specific activity patterns. Unexpectedly, reorientation of larval head involves muscle tone in the apparently motionless posterior segments. With a STGCN(spatial temporal graph convolution neural network)-Generator model, sequence of larval behavioral poses outlined by morphological patterns of muscles could be accurately predicted based on the time series of ventral but not dorsal muscle activities, and vice versa. Laser ablation of ventral but not dorsal muscles interrupted peristaltic wave and undermined head cast in both frequency and amplitude. Our results provide a simplified muscle activity representation of soft body motion that can be used for probing the key components of animal motor control.

## Introduction

Animal behaviors are implemented through the coordinated contraction and extension of its body muscles under the control of motor nervous system. To get a full understanding of neural coding of behaviors, how neural signals are encoded in activity pattern of muscle system and how muscle activity is transformed into final motor output need to be known. Compared with the large body of studies on neural representation of behaviors and muscle activities^1, 2^, the muscle activity representation of behaviors at whole body level is largely lacking. In most animals, to obtain an overall muscle activity representation of body movements at single muscle resolution at high enough spatiotemporal resolution is technically daunting: it needs to monitor the activity of all the related muscles in spatial scope varying from as big as the whole body to as small as individual muscles in freely behaving animals, which is usually out of the capability range of current technologies including electrophysiological recording and optic imaging. Nevertheless, in small and transparent animals such as *C. elegans*, it is possible to monitor the activity of individual neurons and muscles and the overall behavior under a microscope^3, 4^.

The model animal *Drosophila* larva is also small and transparent, making it suitable for simultaneous tracking of behavior and activity of neurons and muscles^5, 6^. It possesses three thoracic segments (T1-T3) and nine abdominal segments (A1-A9), with each semi-segment containing 30 body wall muscles in segments A1-A7 that can be visually identified under microscope^6^. *Drosophila* larva has a rich reservoir of behavioral movement^7^. In recent years, much work has been done to unravel the neural mechanism underpinning cardinal movements such as forward and backward peristaltic crawling at single neuron resolution^8–11^, left-right balance^12^, bending^13, 14^, up-righting^15^, rolling^16^ and so on. The action of body wall muscles has also been closely observed especially during peristalsis^8^. The expression of fluorescent proteins in muscles allowed observation of details of movement and deformation of larval segments at single muscle level^17–19^. In a recent report, calcium imaging has also been used to monitor the activity of all muscles at single muscle level in one side of A1 and A2 segment during forward and backward peristalsis under a confocal microscope at high temporal resolution^6^. The underlying neural activity pattern was elucidated based on neuromuscular connectivity and the observed temporal profiles of muscle activities.

Here in this work, we set up a light sheet system in combination with calcium imaging to monitor the activity of muscle system in a nearly freely behaving 1^st^ or early 2^nd^ instar *Drosophila* larva. We investigated muscle behaviors at single muscle resolution and described dorsal and ventral muscle activity representation of larval forward/backward peristalsis, head cast as well as turning. We found that muscle activity could be highly correlated with morphological properties such as length and orientation angle, especially for ventral muscles. Furthermore, larval pose sequence could be well predicted based on ventral but not dorsal muscle activity sequences using STGCN-Generator model, and vice versa. The requirement of ventral muscles for larval peristalsis and head cast was confirmed by laser ablation of ventral muscles. The establishment of the quantitative relationship between muscle activity pattern and pose sequence will greatly facilitate our understanding of motor control in *Drosophila* larval behaviors.

## Results

### 1. Calcium imaging of larval muscle activity using light sheet microscope

We set up a light sheet imaging system to monitor the activity of the muscles across larval body^20^ (Figure 1a, Figure 1-figure supplement 1). 1^st^ or early 2^nd^ instar larvae expressing activity indicator GCAMP7.0^21^ driven by a muscle specific driver R44H10-LexA^6^ was used. The selective plane illumination of light sheet microscope allowed us to capture quick movements such as mouth extension-retraction and the temporal details of regular locomotion and body bending movements, at high spatiotemporal resolution (Figure 1b and 1c, Video 1 and 2). As larva body was thick, we flipped the chip containing the larva (see Methods for details) bottom-up to obtain clear images of the ventral muscles (Figure 1a). We chose to quantify the properties of dorsal muscle 9 and 10 and ventral oblique muscle 15 and16 throughout whole larval body, since they could be readily recognized (Figure 1d and 1e, Video 3 and 4). Additional dorsal muscles in A8 segment and ventral muscles in T2 and T3 segment were not used in behavioral analysis for simplicity but were used for pose prediction in the following. These properties included calcium signal intensity, length and orientation angle of each muscle (see supplementary File 1A for definition of parameters used in this and following figures). To ensure that calcium signal truly reflected muscle activity^18^, we compared behavior of muscle 9 in segment A1 expressing calcium sensitive GCAMP and calcium insensitive GFP in forward peristalsis. Upon similar extent of muscle contraction, peak increase in GCAMP signal was about ten times that of GFP signal (Figure 1f and 1g). This meant that the GCAMP signal change originated mostly from muscle activity but not muscle deformation. Furthermore, increase of fluorescent intensity of GCAMP happened before muscle contraction (Figure 1-figure supplement 2). The time delay in muscle 9 in different body segments in peristaltic larvae were generally more than 200ms on average. As GCAMP indicated calcium signal could represent the activity of the muscles, we directly used the calcium imaging signal to measure muscle activity in the following experiments.

**Figure 1.**
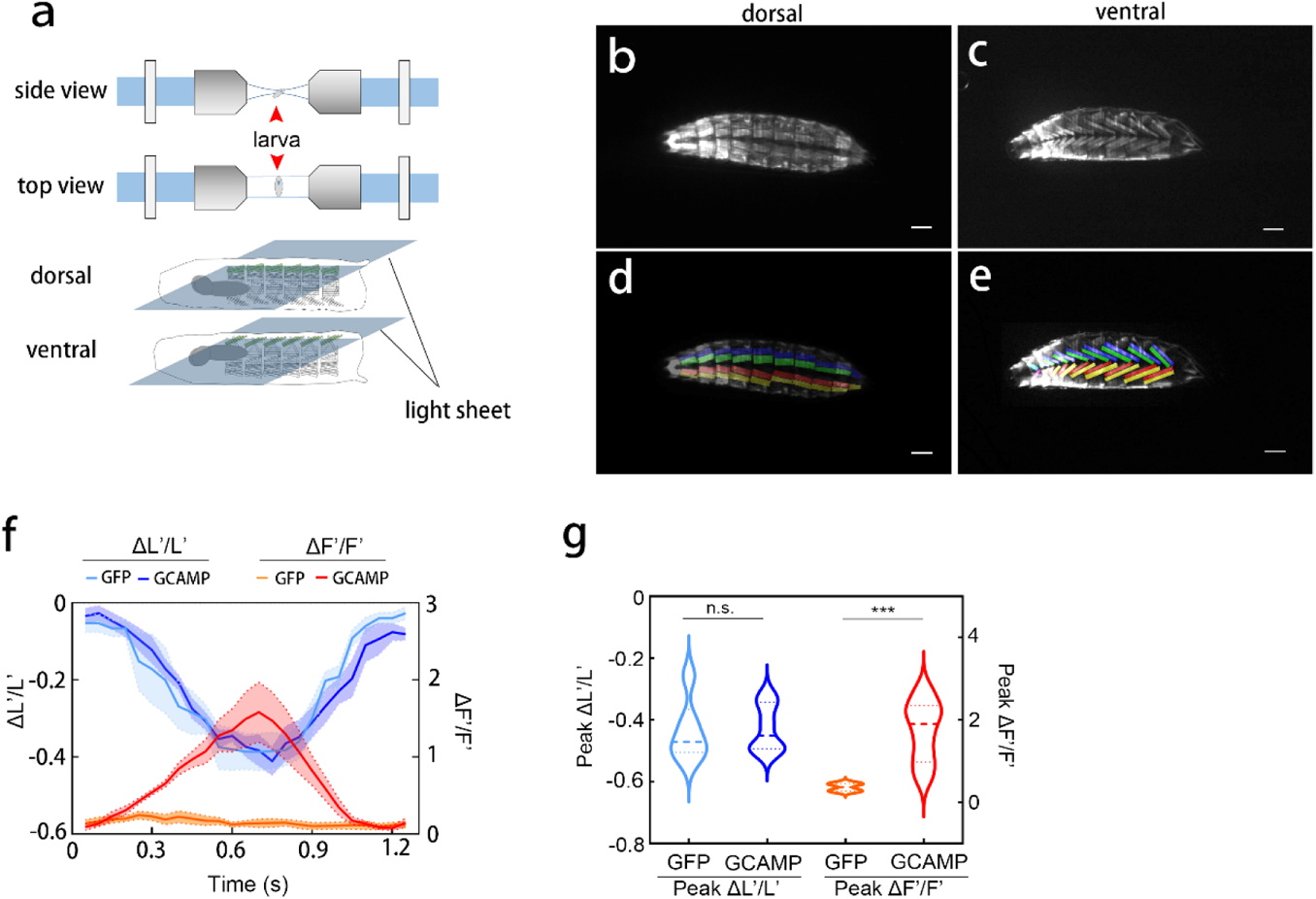
Light-sheet imaging of *Drosophila* larval muscle activities. a. Scheme of light sheet imaging of *Drosophila* larva.
b-e. Imaging and labeling of larval dorsal (b, d) and ventral (c, e) muscles. Larval genotype is *R44H10>GCAMP7.0*. Scale bars, 100um.
f. Curves of activity and length of muscle 9 expressing GCAMP or GFP. Shaded areas flanking curves represented SEM.
g. Quantification of peak muscle activity and maximal contraction in f. Maximal muscle activity change and maximal contraction are respectively indicated by peak ΔF’/F’ and peak ΔL’/L’. n = 6 for each. n.s. not significant, \*\*\**P*<0.001, Mann-Whitney test. Thick line and dot-line in violin plot, median and interquartile range.

**Figure 2.**
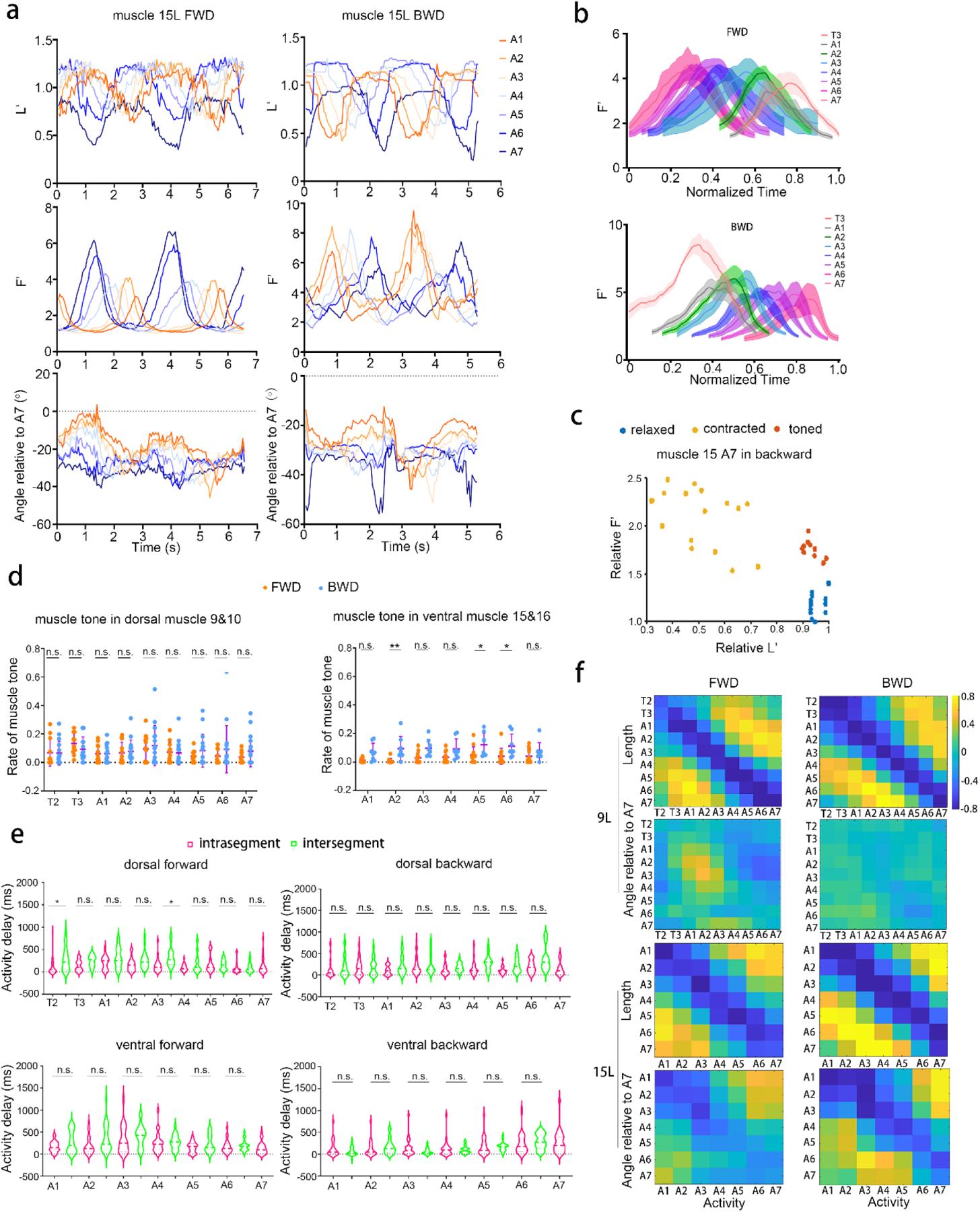
Muscle activity pattern representing larval peristalsis. a. Activity, length and orientation angle of representative ventral muscle 15 during two waves of forward and backward peristalsis.
b. Average normalized one-wave activity pattern of dorsal muscle 9 on left in forward and backward peristalsis. n = 6 for forward; n = 5 for backward.
c. Scatter plot of activity and length of a ventral muscle 15 in A7 segment during a wave of backward peristalsis. The muscle states can be relaxed, contracted and toned, as indicated by blue, orange and brown dots.
d. Rate of muscle tone in the whole peristaltic periods in dorsal (left) and ventral (right) muscles during forward and backward peristalsis. While rate of muscle tone in dorsal muscles during backward is similar to that during forward peristalsis, rate of muscle tone in ventral muscles during backward peristalsis is generally higher than in forward peristalsis. n.s. not significant. \**P*<0.05, \*\**P*<0.01, Sidak’s multiple comparison test after two-way ANOVA. Error bars, SD.
e. Inter-segmental and intra-segmental muscle activity delay in dorsal and ventral muscles in forward and backward peristalsis. The delays within one segment and between neighboring segments are generally not significantly different. Inter-segmental delays and intra-segmental delays are shown in green and magenta respectively. The labels for intra-segment that is supposed to lie between two neighboring segments are not shown due to limited space. n.s. not significant, * *P*<0.05, Sidak’s multiple comparison test after two-way ANOVA. n = 24 for all cases. Thick line and dot-line in violin plot, median and interquartile range.
f. Average correlation between muscle activity and muscle length and orientation angle. Muscle length is strongly negatively correlated with the activity of the same muscle in both dorsal and ventral muscles in forward and backward crawling. Negative correlation between orientation angle and activity of the same muscle is clear in ventral but not in dorsal muscles for both forward and backward peristalsis. The strong positive correlation between activity of muscle in one segment and length and angle of the muscles about four segments away is due to phase delay in peristaltic wave. For dorsal muscle 9, n = 8 waves for forward and n = 6 for backward; for ventral muscle 15, n = 6 for forward and n = 4 for backward.

We thus used the three fundamental parameters of calcium signal intensity, length and orientation angle to describe muscle behaviors during forward/backward peristalsis and re-orientation maneuvers including head cast and turning. We described the temporal properties of muscle behaviors and related larval body pose changes in details as shown in the following. Importantly, the correlation between muscle activity and parameters that describes larval pose was also investigated to disclose the possible relationships. For these behavioral modes, both dorsal muscles (number 9 and 10) and ventral muscles (number 15 and 16) were analyzed.

### 2. Behaviors of muscles in the same segment were highly asynchronized at segmental level in peristalsis

We first looked the larval muscle activity patterns during forward and backward peristalsis. Larval tailspeed was in the range of 110-230 μm/s in forward crawling and 200-300 μm/s in backward crawling. As shown in Figure 2a and Figure 2-figure supplement 1, muscles were activated largely serially from posterior segment to anterior segment to form a forward-propagating wave paralleling the forward wave of the muscle length oscillation in forward crawling. The direction of propagation waves was reversed in backward crawling. Similar temporal sequence in muscle orientation angle was seen in ventral muscles but not as clear in dorsal muscles. As the orientation of larval tail was stable in peristalsis, the relative angle of muscle with respect to the midline of the posterior A7 segment in larval tail was used here to measure muscle orientation. We next looked at the temporal patterns of the muscle activity waves throughout the whole larval body.

As expected, the activation of the segmentally homologous muscles (muscles with same number IDs) on the same side (left or right) occurred largely sequentially in the direction of peristalsis (Figure 2b, Figure 2-figure supplement 2). However, the temporal pattern in backward peristalsis was not the simple reversion of that in forward peristalsis. For certain muscles in backward peristalsis, muscle activation in A1 and A2 segments was largely simultaneous while in forward crawling muscle activation in A2 was always earlier than in A1 segment. Interestingly, the curves of ventral muscle activity waves in backward peristalsis seemed to be skewed rightward in later-active muscles, indicative of slower activation and faster inactivation. This was interesting because the early phase of increase in activity usually was accompanied with no obvious change in muscle length. This corresponded to a muscle tone like state like in human skeletal muscles^22–24^ (Video 5). As shown in Figure 2c, muscles could be in three arbitrarily defined states of contracted, relaxed and toned (see supplementary File 1A for details of definition). In contracted state, muscle was active and contracted. In relaxed state, muscle was completely quiet and fully relaxed. In toned state, muscle was active but did not obviously contract. Muscle tone usually occupied on average less than 10% of the time, but in some cases the occupation rate could be more than 20% in forward and backward peristalsis (Figure 2d). Interestingly, muscle tone occupation rate in dorsal muscles was similar between forward and backward peristalsis whereas the occupation rate in ventral muscles was significantly higher in backward than in forward peristalsis (Figure 2d). This was probably due to the oblique orientation of muscle 15 and 16 that made these muscles differentially prepared for the forward and backward peristalsis.

We next quantified the spatiotemporal pattern of muscle behaviors by calculating the delays of muscle activation and contraction at intersegment and intra-segment level. The mean intersegmental delays between homologous muscles in neighboring segments were all in consistence with the direction of wave propagation, that was, muscles in segments leading the wave generally behaved earlier. This indicated the temporal order was largely well kept at segmental level (Figure 2e, Figure 2-figure supplement 3). We further looked at the intra-segmental delays of muscle behaviors. To our surprise, the absolute value of intra-segmental delays in muscle activation, including those between bilaterally homologous muscles and nonhomologous muscles on the same side, were not significantly different from that of the intersegmental delays involving neighboring segment in most cases (Figure 2e). This meant that temporal range of overall muscle behaviors in one segment overlapped with that in the next segment during peristalsis, although temporal order at segment level had formed in homologous muscles. This was more obvious for ventral muscles since muscle 15 and 16 span two neighboring segments along longitudinal axis. Such temporal overlapping of muscle behaviors along larval body segments obviously facilitated the spatiotemporal continuity and smoothness of larval peristalsis.

**Figure 3.**
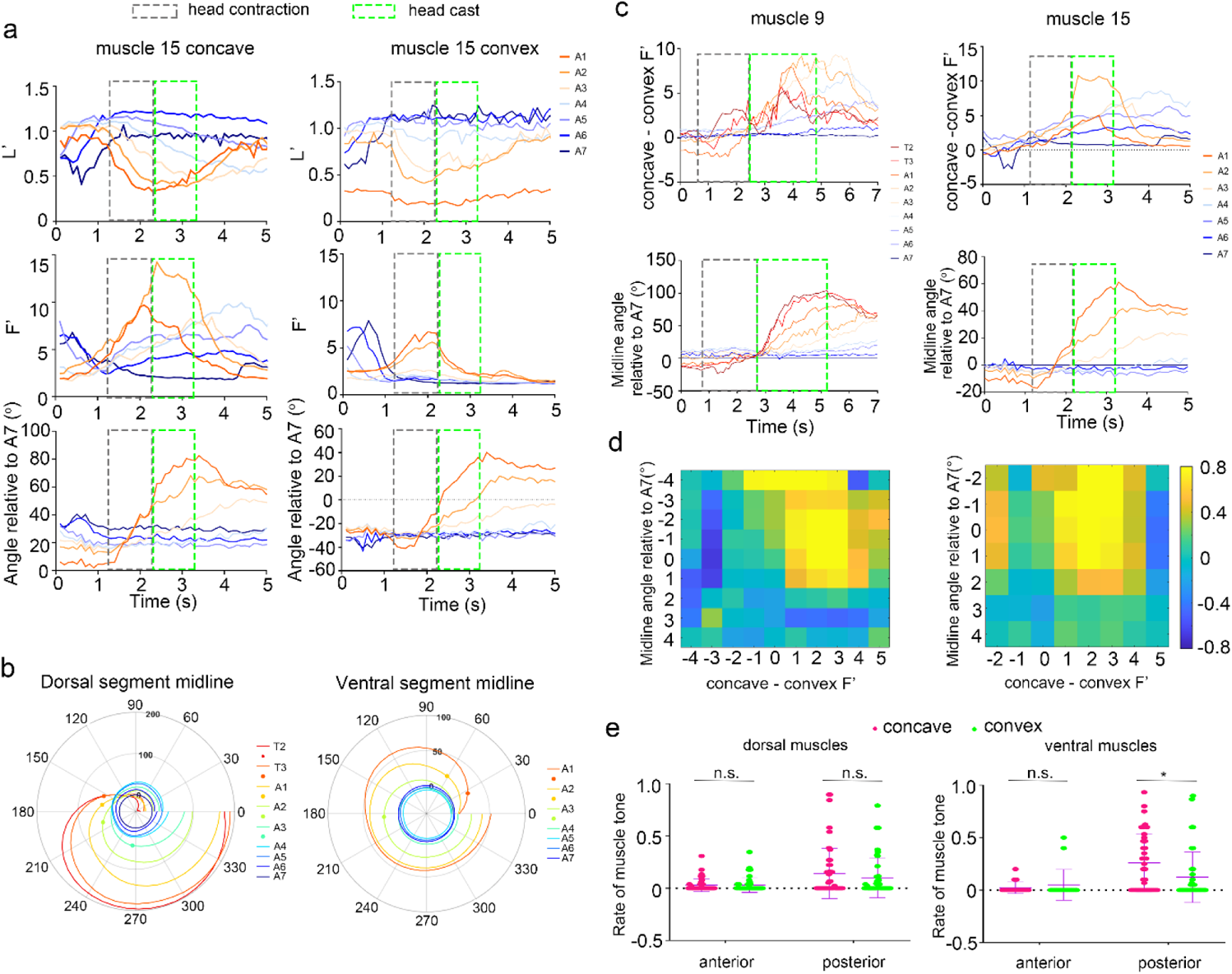
Muscle activity representation of larval head cast. a. Representative curves of activity, length and orientation angle change of ventral muscle 15 on concave and convex side during larval head cast. The grey dotted boxes represent periods of head contraction. The green dotted boxed represent periods of re-orientation in anterior segments.
b. Polar plot of segment orientation calculated based on dorsal and ventral muscles. Dots mark the times that the segment orientation angles reach threshold level. Radius indicates orientation angle; radian of circles indicates time in the range of 5 seconds after the start of head contraction. Note that the more anterior the segments reach the threshold earlier.
c. Representative curves of activity difference between concave and convex muscles and segment midline orientation calculated based on dorsal muscle 9 and ventral muscle 15 during larval head cast. The grey dotted boxes represent periods of head contraction. The green dotted boxed represent periods of re-orientation in anterior segments. Note that during head cast stage, the curves of activity difference in A3, A4 and A5 segment, and curves of segment orientation in segments anterior to A4, are monotonically increasing. Bending points are generally around A2.
d. Mean correlation between concave-convex muscle activity difference and segment orientation angle calculated based on muscle 9 (left) and muscle 15 (right). Segment number 0 is the bending segment. Negative and positive numbers indicate segments anterior and posterior to the bending segment respectively. Strong positive correlation is seen between orientation angle of segments anterior to bending point and bilateral activity difference in segments posterior to bending point. n=6 for dorsal muscle 9, n = 5 for ventral muscle 15.
e. Rate of muscle tone in dorsal and ventral muscles on concave and convex side during head cast. Note that in segments posterior to bending segment, rate of muscle tone is generally higher in ventral muscles on concave side than those on convex side. n.s. not significant, \**P*<0.05, Sidak’s multiple comparison test after two-way ANOVA.

Since muscle activity represented the muscle force that drove larval behavior, we wondered if muscle calcium signal was correlated with muscle length and orientation angle that represented behavioral pose. Indeed, for each type of muscles on both dorsal and ventral sides, strongest negative correlation was mainly observed between activity and length of the same muscle, suggesting that the deformation of muscle was most likely to be caused by the activity change in that muscle itself although the contribution from other muscles could not be excluded. However, obvious correlation between activity and orientation angle was only seen in ventral but not dorsal muscles (Figure 2f, Figure 2-figure supplement 4). This was probably because ventral muscles are obliquely positioned so that their orientation angles are more sensitive to longitudinal contraction than that of dorsal muscles.

**Figure 4.**
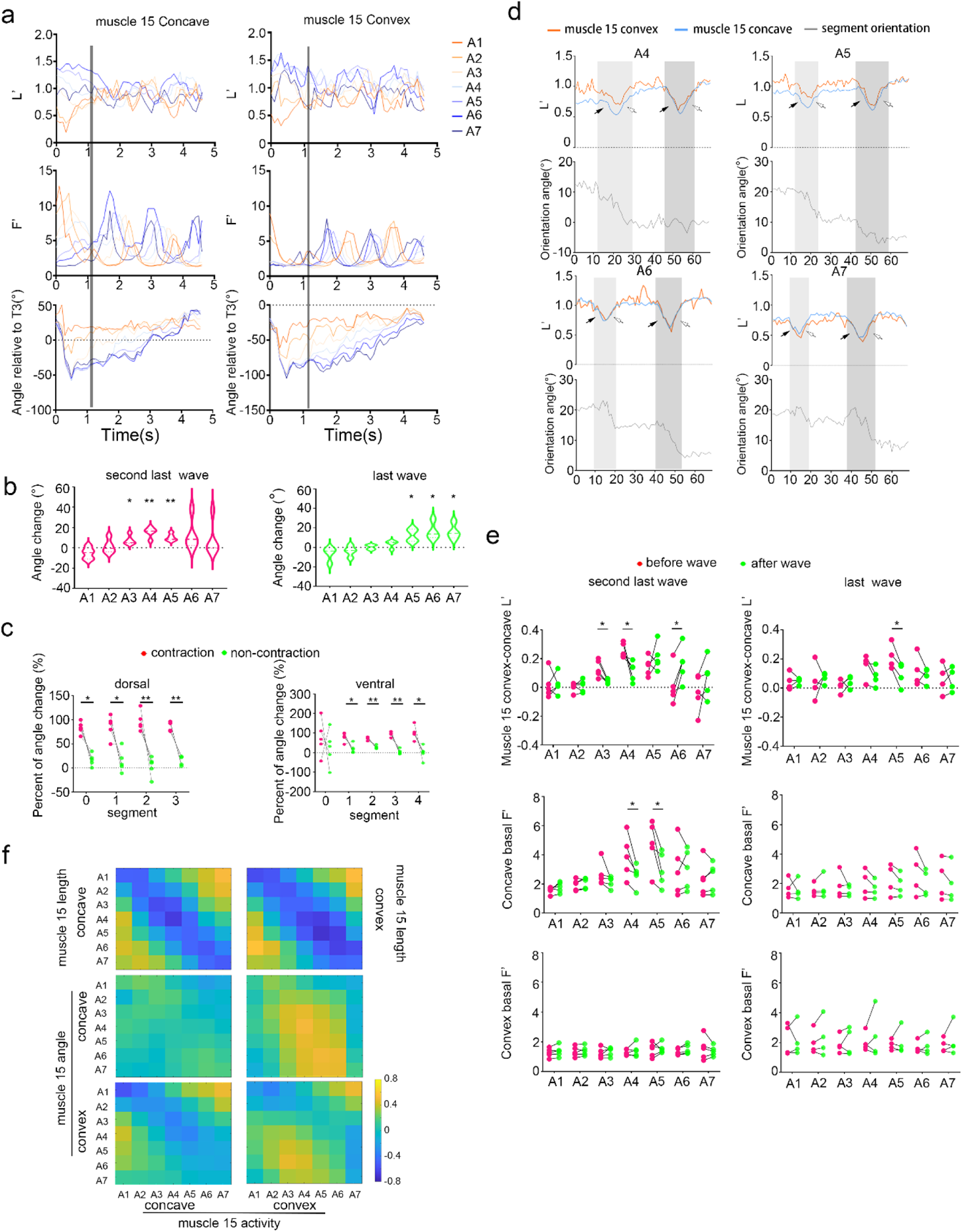
muscle activity representation of larval turning after head cast. a. Representative curves of activity, length and orientation angle change of ventral muscle 15 on concave and convex side during larval turning. The grey lines represent the start of turning after head cast.
b. Segmental orientation changes during second last and last wave of peristalsis in turning before complete alignment of body segments. n = 4 for the last wave, n = 5 for the second last wave. \**P*<0.05, \*\**P*<0.01, one sample *t*-test.
c. Percent of angle change in segments posterior to turning point during peristalsis measured by activities of dorsal (left) and ventral (right) muscles. Segment number 0 is the turning point. Positive numbers indicate segments posterior to the turning. Most angle change happens during crawling stages. \**P*<0.05, \*\**P*<0.01, paired *t*-test.
d. Representative curves of concave and convex muscle 15 length and segment orientation angle in segment A4 to A7 during the last two waves of peristalsis in larval turning. Segment orientation angle is the original angle plus 20° so that the orientation angle of head is around zero (see supplementary File 1A). Solid arrows indicate muscle shortening periods; empty arrows indicate muscle lengthening periods. The dark and light grey areas represent periods of the last waves and the second last wave respectively. Anterior segment re-orientation in A4 and A5 happens mainly in the earlier second last wave, while the last wave mainly changes the orientation of posterior A6 and A7 segment. Note that the change of orientation angle in different segments largely happens in the same period. In more anterior segments, the reorientation happens during muscle shortening and lengthening. In posterior segments, the reorientation happens mainly during muscle lengthening.
e. Change of bilateral length difference in muscle 15 (top row), activity of muscle 15 on concave (middle row) and convex side (bottom row) after the last and second last peristaltic wave during turning. The turning point is at around A3 before the second last wave and around A5 before the last wave. * *P*<0.05, ** *P* < 0. 01, *** *P* < 0.001, paired *t*-test.
f. Mean correlation between muscle activity and length and orientation angle of ventral muscle 15 on concave and convex side. Strong negative correlation is seen between activity and length of the same muscles (top row). Correlation pattern is seen between convex muscle activity and orientation angle of muscles on both sides (middle and bottom row). n = 5.

Thus, in the apparently temporally organized larval peristalsis, the intra-segmental asynchrony in muscles behaviors could contribute to the spatiotemporal continuity of larval peristaltic movements. Larval peristalsis was likely more closely related to ventral muscles than to dorsal muscles.

### 3. Muscle activity was bilaterally asymmetrical in segments posterior to bending point during head cast

While peristalsis relocates larva, head cast and turning are two other fundamental behavioral maneuvers that change larval locomotion direction. We first focused on head cast proceeding forward peristalsis here (Video 6).

During head cast, only larval anterior segments reoriented while the posterior segments were largely quiet (Figure 3a, Figure 3-figure supplement 1). We used the relative angles of muscles or segments with respect to the midline of the posterior A7 segment in larval tail as measurement of orientation. Before the change of orientation angle, larval anterior segments in the head part always contracted, as shown in Figure 3a and Figure 3-figure supplement 1. The orientation angles of anterior segments began to increase before maximal muscle contraction and peak activity were reached in these segments. We thus arbitrarily defined the period between the time that concave muscles in T3 segment began to contract and the time that T3 segment orientation angle relative to A7 reached 20° as the stage of head contraction. 20° was arbitrarily chosed as threshold because it marks the least obvious bending. The following time before reaching maximal T3 segment orientation angle was defined as the stage of head cast. It should be noted that for ventral view data the ventral oblique muscle VO2 in T3 segment was used instead of ventral muscle 15 and 16, since the latters are absent in T3 segment. In head contraction stage, muscles in anterior segments, usually from T1 to A1, were active and contracted. In the head cast stage, muscle activities in these segments decreased and muscles elongated, along with the orientation angles increasing sequentially from anterior to posterior (Figure 3b). The most posterior segment that crossed the re-orientation threshold of 20° was defined as bending point.

We wondered if larval re-orientation was correlated with muscle activities. Plot of muscle activity against muscle length and angle during head cast stage revealed no obvious correlation between length and acitivity for both dorsal and ventral muscles. However, positive correlation could be seen between muscle angle in segments anterior to bending point and muscle activity in segments posterior to bending point (usually at A2 or A3) for ventral muscles and dorsal muscle 10 on concave but not convex side (Figure 3-figure supplement 2). Since reorientation was left-right asymmetric, we then asked if there existed left-right asymmetry in muscle activity. We plotted the time series of concave-convex muscle activity difference and compared it with that of segment orientation angle. Both the curves of orientation angle in anterior segments and that of muscle activity difference in posterior segments were monotonically increasing in the stage of head cast (Figure 3c). Spearman’s correlation analysis showed that concave-convex muscle activity difference in the segments posterior to bending point were highly correlated with the angles of the segments anterior to bending point, for both dorsal and ventral muscles (Figure 3d, Figure 3-figure supplement 3). This suggested that muscles in posterior segments might contributed to the reorientation of anterior segments.

We then looked closely at muscle activity in segments posterior to bending point. While the muscles on convex side in many cases were simply quiet without obvious change in activity and length, muscles on concave sides were always active (such as concave and convex muscle 15 in A4 and A5 segment in Figure 3a and Figure 3-figure supplement 1, Video 7). The concave muscles in segments immediately posterior to bending point generally contracted slightly. The ratio of muscle length to activity was generally lower in concave than in convex muscles in segments immediately posterior to bending point, suggesting that concave muscles were more apt to be active without contraction compared with convex muscles (Figure 3-figure supplement 4). In consistence with this, muscle tone was obviously more frequent on concave side than on convex side in ventral muscles, although such asymmetry was not seen in dorsal muscles (Figure 3e). Thus these muscle tone type of ventral muscle behavior in post-bending-point segments on concave side were closely related to head cast. We hypothesized that the post-bending-point segment muscles contributed to head cast at least in two ways: first, the slight muscle contraction on concave side not only provided part of the pulling force on anterior segments but also saved necessary space for head cast towards concave side; second, muscle tone probably changed the stiffness of muscles and thus could provide stronger support for the moving anterior segments. It was noted that contribution of other muscles to head cast should not be ignored.

In short, the reorientation of anterior segments in larval head cast involved the seemingly quiet posterior segments.

### 4. Turning was signaled by ventral muscles but not dorsal muscles

After the head cast, a larva might change its orientation by aligning the posterior segments with the new direction of head. For simplicity, we focused only on the cases in which orientation of head was largely stable after head cast. Different from in peristalsis and head cast, all orientation angles were calculated with respect to that of T3 segment to demonstrate the process of body alignment towards head since the orientation angle of larval head was relatively stable (Figure 4a, Figure 4-figure supplement 1, supplementary File1A). The most anterior segment with absolute value of relative orientation angle above 20° was defined as turning point, similar to the bending point in head cast. The re-alignment of posterior segments with anterior segment, or turning, was accomplished by forward peristalsis. As shown in Figure 4a and Figure 4-figure supplement 1, while muscles in all segments were shortened and lengthened periodically, orientation angles in segments posterior to turning point decreased progressively along with the progress of the peristaltic wave. In our experiments, one or two waves of peristalsis were generally required for the larva to completely realign posterior segments with head. In the earlier second last wave, larva mainly adjusted orientation of segments closely posterior to the turning point (usually at A2 or A3) to align with the anterior part (Figure 4b). After the wave, the turning point was shifted posteriorly usually to A4. In the last wave, larva mainly adjusted orientation of posterior segments such as A5 to A7 (Figure 4b). It should be noted thatwe used the original angles but not the relative angles of the segments to exclude the effect of head movement on relative segment orientation angle (see supplementary File 1A). Therefore, larva adjusted its body segment orientation mainly in the three segments posterior to the turning point in both waves. Most or even all these changes in segment orientation happened in the peristaltic periods that the segments were contracted, no matter in dorsal view or ventral view (Figure 4c).

We next asked how segment reorientation was correlated with muscle contraction. As shown in Figure 4d, for ventral muscles in segments posterior to turning point, more of the reorientation happened during muscle shortening in the more anterior segments, while in more posterior segments more of the reorientation happened during muscle lengthening. As the lengthening of posterior segments happened concurrently with the shortening of anterior segments during the forward propagation of peristaltic wave, the reorientation of these segments happened largely at the same time, that was, when the segment close to turning point was being shortened. It was thus likely that orientation change was caused by the contraction of the segments around the turning point. We noticed that after the peristaltic waves, the length of concave muscles was increased in segments close to the turning point so as to reduce the bilateral muscle length difference that reflected the extent of body bending (Figure 4e). Accordingly, the high basal activity levels of concave muscles were reduced after peristaltic wave while that of convex muscles remained stable at relatively low level (Figure 4e). The changes in the last wave in most cases were not significant probably because the bending angles were relatively small (Figure 4e). In dorsal muscles, the change in muscle length asymmetry and basal activity level was not obvious, although segment reorientation was still clear (Figure 4-figure supplement 2). Thus at least for ventral muscles, peristaltic wave realigned the bilateral muscles not only in length but also in activity level. Such bilateral symmetry promoting realignment was likely the driving force of straightening of larval body during turning. One advantage of realization of turning through peristalsis is probably that larva could change its body orientation and crawl forward at the same time.

As turning involved asymmetric peristalsis, we asked if the correlation between muscle activity and muscle length and angle could still be seen. While the negative correlation between activity and length of the same muscles was seen like in normal peristalsis (Figure 4f, Figure 4-figure supplement 3), the correlation between activity and angle in ventral muscles was no longer as clear as in peristalsis (Figure 4f, Figure 4-figure supplement 4). We then widened our scope of correlation from between homologous muscles on the same side to between all concave and convex muscles. We found that orientation angles of muscles in segments posterior to turning segment (mostly A3) on both concave and convex side seemed to be positively correlated with the activity of convex muscles in segments posterior to turning point (except for A7 segment) for ventral muscles (Figure 4f). On the other hand, no obvious correlation pattern was found for dorsal muscles (Figure 4-figure supplement 4). The difference in angle-activity correlation in convex and concave muscles was probably caused by some unknown bilateral temporal difference.

Therefore, ventral muscles also out performed dorsal muscles in representing turning behavior.

### 5. Larval pose sequence could be predicted based on ventral muscle activity patterns

Since larval behavioral movement was the consequence of muscle activity, muscle activity patterns should contain enough information to generate the corresponding larval pose sequences. As shown above, muscle length and angles could be well correlated with muscle activity patterns in some cases for dorsal muscle 9/10 and ventral muscle 15/16. Since only limited number of correlations were investigated, there could be many other correlations that had not been discovered. We postulated that muscle activity pattern might be able to represent larval pose which contains information not only including length and orientation angle of each muscle but also the whole spatial pattern of the muscles. We should be able to predict larval pose sequence using muscle activity data.

By focusing on activity of each individual muscle and overall pose represented by the set of muscles, we developed a STGCN generator model consisting of graph convolutional neural network (GCN) and Long-Short term memory (LSTM) to translate the time series of muscle activity into temporal sequences of muscle morphological properties (length, width, angle, and center coordinates of muscle 9 and 10 in segment T2-A7 plus one pair of similar muscles in A8 segment on dorsal side, and of muscle 15 and 16 in segment A1-A7 plus two pairs of similar muscles in T2 and T3 segment on ventral side), as shown in Figure 5a. The following two facts had been considered in this model: first, the current pose depends on the previous pose and muscle activities; second, the relationships between muscle activities and pose sequences is general^25, 26^ (see supplementary File 1B for details). Given an arbitrary starting pose and subsequent muscle activity sequence in behavioral modes including peristalsis, head cast and turning, the model was able to generate subsequent pose sequences that resemble true pose sequences of more than 6 seconds long, after training with 2000 epochs on observed ventral and dorsal muscle imaging data (Figure 5b). The disparities calculated by Procrustes analysis between the generated and the ground-truth larval pose was also shown. When pose disparities were less than 0.02, the model generated poses were indistinguishable from the realistic ones. From the pose disparity curve and the sampled prediction-real poses, we could see that the pose sequences generated using ventral muscle data closely approximate the sequences of realistic larval poses (Figure 5b; Video 8-10). On the other hand, the overall pose sequences generated using dorsal muscle data was different from the ground truth (Figure 5b, Video 11-13). As shown in Figure 5c, pose disparity was significantly increased in pose sequences based on dorsal muscle activity patterns compared with those based on ventral muscle activity patterns. Therefore, the model trained on ventral muscle data was more promising for modeling relationships between muscle activity pattern and corresponding larval pose.

**Figure 5.**
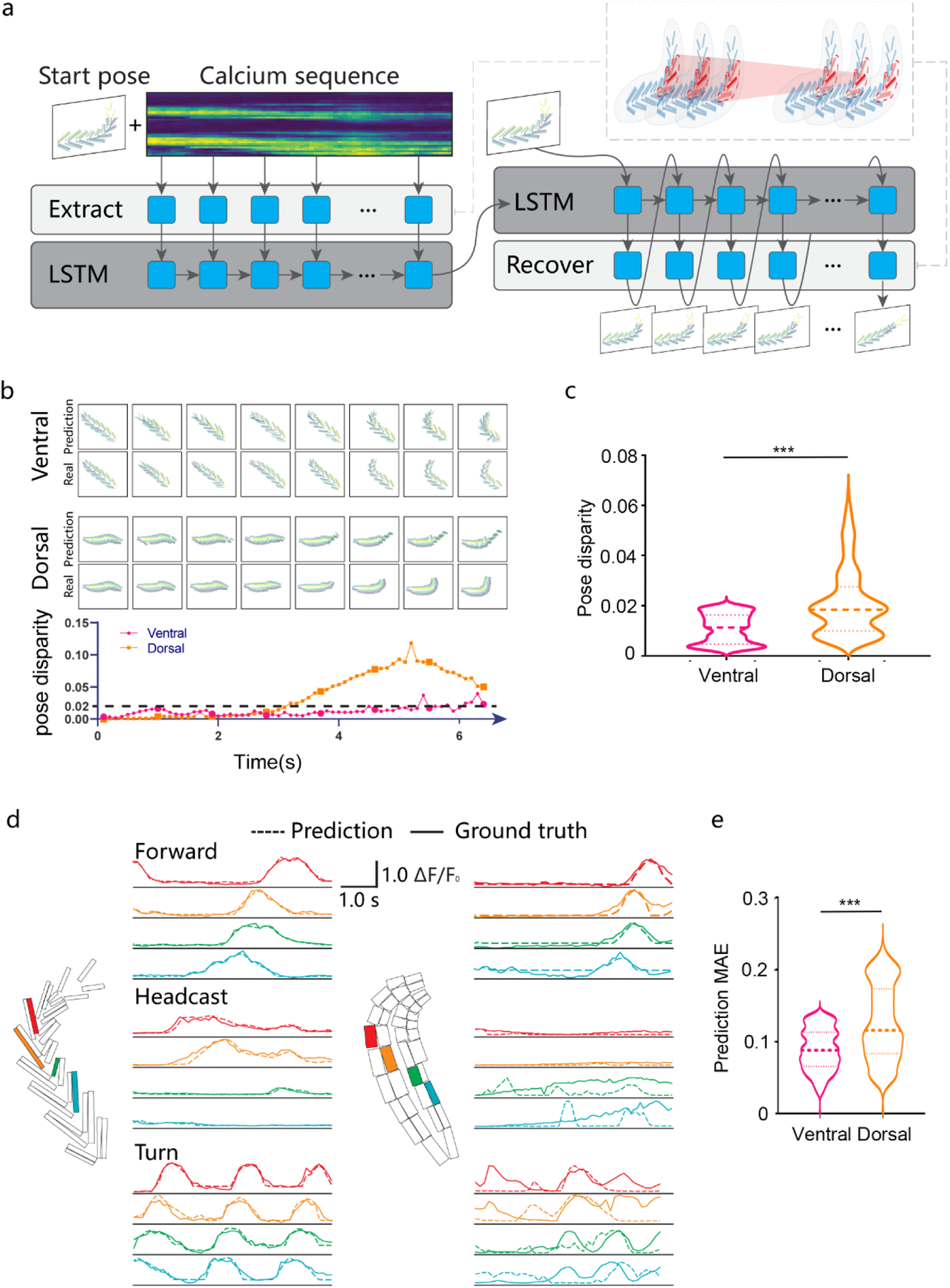
Larval pose sequence can be predicted based on sequences of ventral muscle activities. a. Schematic demonstration of STGCN generator model for end-to-end calcium-muscle behavior translation. The model takes a frame of muscle pose and a set of muscle calcium activity sequences as input, and then generates the subsequent pose sequence following initial muscle pose under the encoding of corresponding muscle calcium activity sequences. The model works in an autoregressive manner, each of the next pose is estimated based on historical poses and muscle activities. And a graph convolutional network is used for fusing this historical information. Example frames of input ventral poses is shown on right side. Each piece of muscle is drawn with blue polygons. The receptive fields of a filter with D = 1 are drawn with red dashed circles for fusing the spatial-temporal information of muscle activities.
b. Pose sequences generated based on activity sequences of dorsal and ventral muscle. The real and predicted pictures in top and middle row, corresponding to the large circle or squares in the bottom panel, are sampled once every 10 frames from the original frame sequence. The pose disparity is calculated by Procrustes analysis. The lower the disparity, the closer the predicted pose is to the ground truth pose.
c. Statistics of overall pose-level muscle behavior disparities. *** *P*<0.001, Mann-Whitney test.. n =244 for dorsal, n = 268 for ventral. Thick line and dot-line in violin plot, median and interquartile range.
d. Muscle calcium activities generated based on pose sequences. Curves of representative dorsal and ventral muscles during forward crawling, head cast and turn are shown.
e. Statistics of overall muscle calcium activities prediction errors. The mean average error (MAE) of the model trained on ventral data is statistically significantly lower than the model trained on dorsal data. \*\*\**P*<0.001, Mann-Whitney test. n = 470 for ventral, n = 249 for dorsal. Thick line and dot-line in violin plot, median and interquartile range.

On the other hand, we wondered if larval muscle activity patterns could be predicted based on larval pose. As pose could be considered as the consequence of past muscle activity, the prediction of muscle activity was performed in a rewinding manner, which means that the muscle activity sequence was generated from end to start based on correspondingly reversed pose sequences. It could be seen that in behavioral modes such as forward peristalsis, head cast and turn, the pose-sequence-based prediction of ventral muscle activities were very close to the true muscle activities, whereas such predicted dorsal muscle activities often deviated away from the true signals (Figure 5d). As depicted in Figure 5e, the prediction of mean average error (MAE) of generated calcium activity sequences was significantly smaller in ventral muscles than in dorsal muscles, i.e., prediction of ventral muscle activities based on pose sequence was more accurate than prediction of dorsal muscle activities.

Taken together, these prediction results were consistent with our observation that the correlation between activity and orientation angle was more significant in ventral muscles than in dorsal muscle. Thus, the activity pattern of the ventral muscle 15 and 16 was able to recapitulate the activity of the whole muscle system that produce larval pose and behavior, although ventral muscles alone should not be enough to generate normal larval behavior.

### 6. Laser ablation of ventral but not dorsal muscles interrupted larval peristalsis

As larval ventral muscle activity pattern could well represent behavioral movements, they might play functional roles in these behaviors. We then tested this hypothesis by ablating these muscles using two photon laser beam^27^ (Figure 6a and 6b). When ventral muscle 15 and 16 in two consecutive segments were ablated bilaterally, spontaneous forward peristalsis started at and backward peristalsis induced by gentle brush touch stopped at the segment of lesion, as if the segments posterior to lesion site were paralyzed. We next bilaterally ablated ventral muscle 15 and 16 in single segment from A1 to A7. Forward and backward peristaltic waves indicated by the rate of change in segment area were found to cut off at, or immediately posterior to, the injured segment, when lesion was in segment A4 or A5 (Figure 6c, Video 14, 15). On the other hand, bilateral laser ablation of dorsal muscle 9 and 10 or skin (used as control) in single segment did not affect the propagation of the peristaltic waves (Figure 6c, Video 14, 16). We quantified the effect of ventral muscle ablation in single segment on completeness of tail-to-head forward peristaltic waves by counting the number of peristaltic waves in T3 and A7 segment that represented head and tail respectively. While ventral muscle lesion in A6 and A7 segment had no effect and those in A1 and A2 segment had occasional interruptive effect on peristaltic wave, such lesion in A3, A4 and A5 segment could always completely interrupt the peristaltic wave (Figure 6d). These observations confirmed that ventral muscles were indeed required for larval peristalsis. On the other hand, ablation of dorsal muscle 9 and 10 bilaterally in single segment did not impair larval peristalsis, suggesting that they were not required for normal crawling (Figure 6d). The effect of muscle lesion on peristaltic wave transmission could also be reflected by the extent of segment contraction in segments neighboring the injured segment. As shown in Figure 6e, contraction in two segments posterior to the lesion site in ventrally injured larvae was not as significant as in the anterior segments, while this was not seen in control larvae and larvae subjected to dorsal muscle lesion. This result further support the function of ventral muscles in larval peristalsis. We also checked impact of muscle ablation on larval head cast. Bilateral ablation of ventral muscle 15 and 16 in segment A3 to A5 significantly decreased the frequency of larval head cast induced by gentle touch in the head while bilateral ablation of dorsal muscle 9 and 10 did not (Figure 6f). Furthermore, the size of head cast was also undermined by ablation of ventral muscles in A3 to A7 but not by that of dorsal muscles in any segment (Figure 6g). This was also consistent with the result of model prediction on ability of ventral muscles in representing larval behaviors such as head cast.

**Figure 6.**
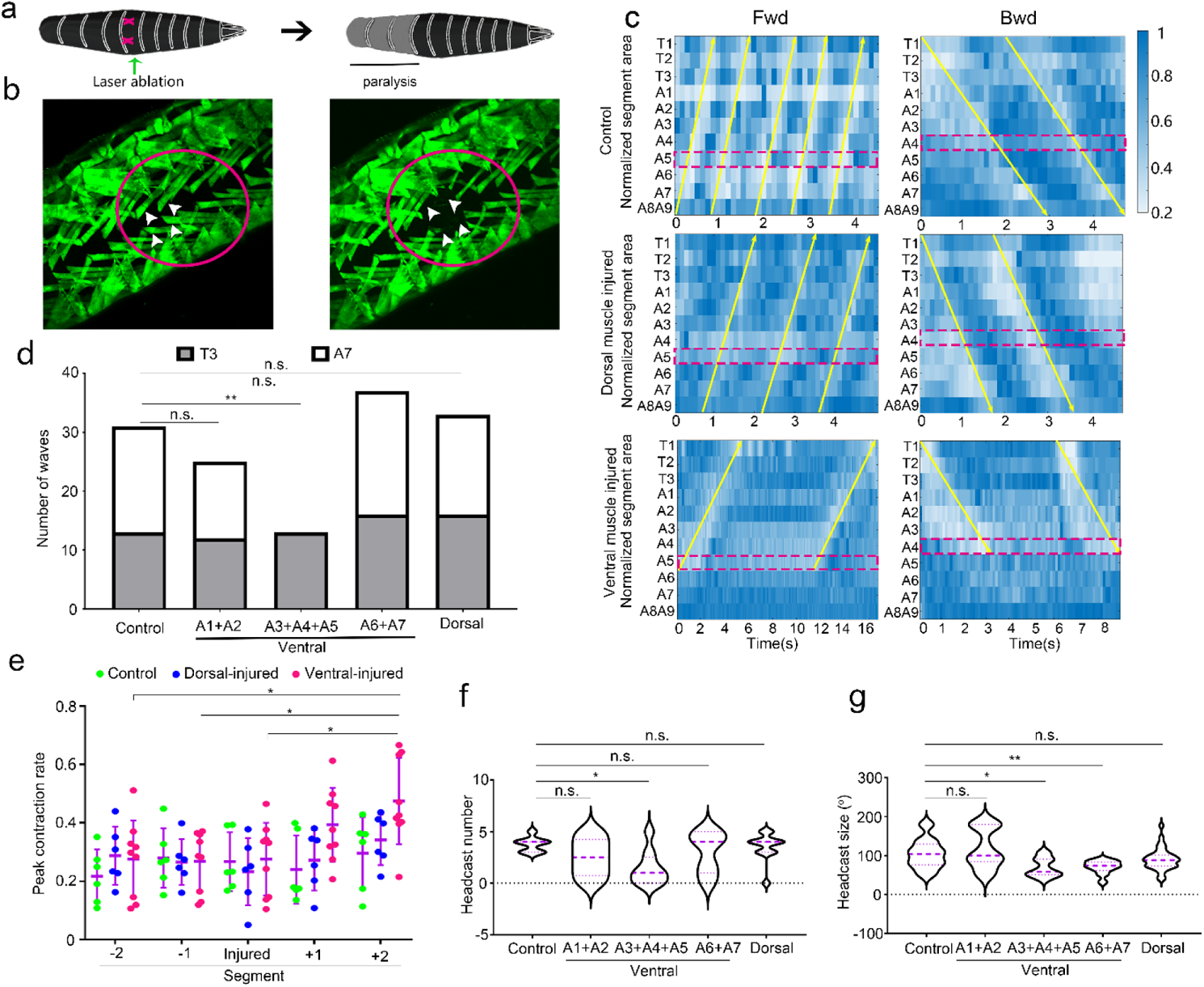
Ablation of ventral but not dorsal muscles in larval undermines larval crawling. a. Schematic representation of the effects of muscle ablation on peristalsis. The magenta crosses indicate the bilateral muscle ablation. Grey blocks indicate paralyzed segments. Black blocks indicate intact segments. When the ventral muscles 15 and 16 are ablated, the segments posterior to the injury are paralyzed.
b. Confocal images of muscle 15 and 16 before and after laser ablation. The positions of muscle 15 and 16 are indicated by arrow heads.
c. Bilateral laser ablation of the ventral muscles 15 and 16, but not dorsal muscle 9 and 10, interrupts both forward and backward peristaltic waves. The long yellow arrows indicate the peristaltic waves in segment area. The magenta dotted rectangles mark the segment of muscle ablation. In control, only skin but not muscles are injured. Note the absence of peristaltic wave in segments posterior to injury in ventral muscle ablation.
d. Comparison of peristaltic wave numbers seen in anterior T3 and posterior A7 segments in larvae subjected to muscle ablation at different segments within two minutes after injury. Bilateral ablation of ventral muscles in A3 to A5 segment completely abolished the peristaltic wave in A7 segment. Ablation of dorsal muscle in all other segments has no obvious effect on completeness of wave propagation.
e. The maximal contraction during peristaltic wave, measured by the ratio of minimal segment area to full segment area, is reduced in segments posterior to ventral muscle ablation. Segment contraction anterior and posterior to the injury is not affected by ablation of dorsal muscles. Injured segment is the segment of muscle ablation. Positive and negative numbers indicate segments posterior and anterior to the injury.
f. Ablation of ventral but not dorsal muscles reduces the frequency of larval head cast. Number of head cast is counted after five times repeated touch in the head for each larva.
g. Ablation of ventral but not dorsal muscles reduces the size of larval head cast. For d-g, n.s. not significant, \**P*<0.05, \*\**P*<0. 01, *Fisher*’s exact test for d, Sidak’s multiple comparison test after two-way ANOVA for e-g. Error bars, SD. Thick line and dot-line in violin plot, median and interquartile range.

Taken together, these results showed that ventral muscles but not dorsal muscles were required for larval peristalsis and head cast, which was in support of the ability of ventral muscles to represent larval behavioral movements.

## Discussion

In this work, we used a self-built light-sheet microscope to acquire the real time panoramic muscle activity pattern accompanying *Drosophila* larval behavioral movements. We found that *Drosophila* larval soft body movements were mediated by intersegmental coordination of muscle activities. At single muscle level, activity of certain muscle was highly correlated with length or orientation of that muscle or other muscles, especially for ventral muscles. Interestingly, ventral but not dorsal muscle activity pattern was sufficient to generate larval pose sequence using a deep neural network model. Furthermore, laser ablation of ventral but not dorsal muscle at segment from A3 to A5 was able to interrupt propagation of peristaltic wave at the segment of lesion and undermine head cast which apparently did not involve the injured segment.

Here we presented a STGCN generator model consisting of graph convolution network and long-short term memory to accomplish the recaption of bidirectional translation between muscle calcium sequence and motor behavior sequence with high accuracy. However, much is still required before we can fully understand such highly complex and synergistic processes. For example, as the data used in this study contain only the calcium activities of a subset of the muscles, other factors such as other muscles that could be involved in pose cannot be excluded. Similarly, although the graph convolution used in this model assumes the spatially closest muscles have the greatest degree of interactions, it is possible that there are muscles controlled by the same source but are spatially far from each other. In spite of these confounders, our model’s capability to produce accurate bidirectional translation over a subset of muscle activities and overall sequence within arbitrary time frames suggests that a subset of ventral muscles is sufficient to generate overall motor behaviors. It is also worth noting that without comprehensive muscle states and external environment feedback, our model is still capable of resolving pose sequence from calcium activities within a group of individual muscles and restoring these calcium activities from the reverse overall posture sequence. Thus, although there may be complex intrinsic muscle dynamics and higher-level control signals that our model fails to account for, this subset of ventral muscles may be sufficient to reveal the dominant patterns of larval motor behavior. Furthermore, our model can potentially be extended to longer sequence translation tasks with lower errors in the future, by using a larger dataset and teacher-forcing techniques.

The finding that larval behavior poses, at lease those observed in the horizontal plane, could be predicted based only on activity of about 30 ventral muscles is especially intriguing. It would necessarily greatly simplify the deciphering of behavioral movement, since we no longer need to analyze the activity information of all ∼600 body muscles to get a full depiction of behavioral poses. It is possible that larval poses could be recapitulated by even fewer muscles. For example, the ventral oblique longitudinal muscle 15 and 16 could be mutually redundant in preserving larval pose since they are not only morphologically similar, but also belong to the similar activation group in peristalsis in A1 segment (Zarin et al., 2019). Furthermore, as each muscle is innervated by specific motor neurons, it is possible to use the activity patterns of these muscles to probe the specific activity patterns of innervating motor neurons or premotor neurons that represent corresponding larval behavioral movements. Reducing the number of representing muscles or neurons can help us quickly get the key factors underlying larval motor control.

Although our results of muscle ablation were in consistence with model prediction, explanation at neural mechanical level is still needed. For disruptive effect of ventral muscle ablation on peristalsis, we assume that spatiotemporal seriality in muscle activation is required for normal larval peristalsis. Since muscle ablation does not injure the central nervous system and the CPG (central pattern generator) that drives peristalsis were intact, the disruption of peristaltic wave by ablation of ventral muscles could be due to loss of feedback from ventral muscles to CPG that is required for further propagation of peristaltic wave. Such feedback signal is probably carried by neurons that sense muscle contraction. Similarly, for effect of muscle ablation on larval head cast, we assume that muscle tone of ventral muscles in posterior segments generate neuronal signals that are required for the activation of the neurons commanding larval head cast, so that ablation of these ventral muscles undermines frequency or size of head cast. These hypotheses need to be further confirmed.

It is unexpected that activities in ventral muscles are more highly correlated with larval behavioral poses than that in dorsal muscles, since dorsal muscle 9 and 10 are apparently the strongest muscles. This is probably because of the following reasons. First, ventral muscles 15 and 16 are obliquely aligned so that their orientation angles are sensitive to peristaltic contractions that are mainly in longitudinal direction. On the contrary, dorsal muscle 9 and 10 are largely parallel to the longitudinal midline so that their orientation angles are insensitive to peristaltic contraction. Second, ventral muscles except for those in A7 segments span two segments which makes them sensitive to deformation in two adjacent segments that include more pose information. Dorsal muscle 9 and 10 however, span only one segment, so that they are relatively less sensitive to deformation in neighboring segment. Third, ventral muscles in normal cases directly touch supporting ground so that they provide the direct supporting force for larval motion. On the other hand, dorsal muscles do not directly exert force on surrounding environment and they provide auxiliary force for larval movement. Although in microfluid chip larva might touch surrounding parts on both dorsal and ventral sides, exerting force with ventral muscles may still be the more natural way.

One of our discovery is the comprehensive occurrence of muscle tone that is equivalent to isometric muscle contraction in mammalian skeletal muscles. In larval peristalsis, muscle tone was frequently seen especially in posterior segments during backward crawling, leading to the rightward skewness of activity curves. In larval head cast, muscle tone in concave ventral muscles always occurs during the reorientation of anterior segments. We propose that in peristalsis muscle tone helps to keep muscles primed and ready for activation, while in head cast muscle tone helps to maintain body balance and postures by changing the stiffness of muscles.

One prominent feature of *Drosophila* larval movement is the soft-style, i.e. continuity and smoothness. Based on our observation, the continuity and smoothness reside not only in the flexibility of the physical structure, but also the spatiotemporal coordination of muscle behaviors. In peristalsis, the temporal range of muscle behaviors in neighboring segments overlaps. Meanwhile, temporal sequence is well established in homologous muscles. Considering that lateral muscles behave later than the longitudinal muscle^6^, the temporal overlap should be even larger. Such spatiotemporal continuity in muscle behaviors supports the apparently continuous but not continual physical movement. The spatiotemporal coordination is also reflected by muscle tone. In peristalsis, muscle tone prepared the muscle for upcoming activation when the wave is still in preceding segments, thus facilitating smoother spatiotemporal transition between segments. In head cast, muscle tone with slight muscle shortening in segments posterior to bending point also allow smoother and coordinated reorientation in anterior segments.

In summary, our work disclosed the activity patterns of reduced number of muscles for describing larval movements and provided simplified physiological signatures for soft body behaviors at single muscle resolution that could facilitate future investigation of animal motor control.

## Materials and methods

### Fly culture and strains

all flies were raised at 25°C on standard medium and 12h:12h light/dark cycles of culture^28^. The following fly strains were used in this work: R44H10-LexA^6^ (BDSC61543), LexAop-GCAMP7s^20^.

### Light-sheet imaging setup

The schematic representation the of light sheet system setup was shown in Figure 1-figure supplement 1. In brief, a 488nm Gaussian laser of 20mw (PL488-20, Shanghai sfolt Inc) was passed through a collimator to form a collimated Gaussian beam with ∼3.3mm diameter light spot, and then a lens followed by a cylindrical lens was used to compress the Gaussian light into 1.65mm in height while keeping the width of light unchanged. The light beam was then passed through a second cylindrical lens to further expand the width of the beam from 3.3mm to 9.9 (1.65mm in height). The reshaped elliptical beam was split into two beam paths using a 50/50 beam splitter, with the first beam sequentially passing an optical slit, a cylindrical lens and an illumination objective to form a thin laser sheet that projected onto the sample; and the second beam passing the similar optical modules except for two additional relay lens that could adjust the 2^nd^ laser sheet to completely align it with the 1^st^ one from the opposite direction.

The fluorescence detection was based on an Olympus MXV10 microscope. To achieve a large enough field of view (FOV), we used a low-magnification 4x objective combined with a Photometrics Iris 15 camera to capture all the tracks of the free movement of a 1^st^ instar or early 2^nd^ larva in a 5mm x 0.6mm square chamber of a customized microfluidic chip. The position and orientation of the sample could be adjusted using a customized sample holder. The light-sheet microscope allowed selective-illuminated, high-contrast fluorescence imaging of the moving muscles at a frame rate of 20 frames per second. Since the larva body was thick (∼120um, the height of the microfluid chip chamber) and induced the signal scattering from the ventral muscles. We then flipped the chip bottom-up to obtain clear images of the ventral muscles. All the recorded images were processed using ImageJ.

### Tracking of muscle signal

The dorsal and ventral muscles were manually tracked by defining the four corner points of each muscle rectangle. The calcium intensity in the rectangle was quantified using a Matlab-based custom script. Note that although we have tried to control the level of light sheet to scan only muscle 9 and 10 on dorsal side, so that the possible partial overlap between areas of muscle 9 and 10 and that of muscle 1 and 2 was minimized. For the rectangle, the middle points between neighboring points were calculated as the average x and y axis values. The longer distance between the opposing middle points were calculated as muscle length while the shorter distance was muscle width. The orientation of the line of muscle length was used as orientation angle of that muscle. The mean orientation of the two bilaterally symmetrical muscles was used as the orientation of each body segment. The orientation angle of segment A7 was subtracted from that of each muscle to obtain the relative orientation angle of each muscle.

For judgment of start of each calcium wave or muscle contraction, the minimal calcium level and maximal muscle length in the whole video that encompass at least two periods of peristalsis was defined as basal calcium level and length of an intact muscle. The time point that calcium signal rise to 10% of basal level was defined as the start time of that calcium wave. The time point that muscle length was reduced to be less than 90% of full length was defined as the start time of that contraction.

### Muscle Laser Ablation

Put a washed clean early 2nd instar larva on a 3% agar plate and cover it with a cover glass. Move the cover glass to roll the larva so that its back or abdomen face the objective of two photon microscope for laser ablation. Under a two-photon microscope (FVMPE-RS, Olympus Inc.) equipped with a 25x objective, find target muscle and place it in the center of field of vision and adjust focus to make the end of muscle to be the clearest. In the software control panel, draw a line at the end of the muscle to be ablated. Line scan the muscle end using an 800nm laser beam at intensity of ∼30% of maximal power for ∼1 second. Compare the image before and after ablation to ensure that the muscle is ablated. After laser ablation, remove the coverslip and transfer the injured larva onto a new 1.5% agar plate. Record larval behaviors on the agar plate. Backward peristalsis and head cast were induced by gentle touch in larval head with a brush. For head cast, each larva was repeatedly touched for five times. The size of head cast was calculated as the maximal angle between midline of segment T3 and the midline of segment A7 midline. Attempted head casts with maximal angles less than 20 degrees were not counted in. The number of head cast after five times of touch was counted for each larva.

### Prediction model

A pair of encoder and decoder constructs an encoder-decoder generator neural network capable of mapping larval pose sequences to muscle activity sequences in a one-to-one manner. The encoder stacks two layers of spatial-temporal graph convolution operators and the LSTM layers. The decoder stacks one LSTM layer, two linear layers and two layers of spatial-temporal graph convolution operators.

The generator neural network is optimized based on the sequence reconstruction loss and adversarial loss. Reconstruction loss is an L1 loss between generated sequence and real sequence. The adversarial loss is introduced by an extra discriminator. The discriminator shares similar architecture of encoder is used to extract the distinguishable features for sequence. This means that the adversarial loss can be summed up as a difference in distribution between generated and real sequences, estimated using a neural network. Finally, we train the discriminator to maximize the adversarial loss, as well as the generator neural network to minimize the reconstruction loss and adversarial loss.

We trained such models from muscle activity to pose and from pose to muscle activity for the dorsal and ventral datasets, respectively. The pose sequence is represented by a tensor **X** ∈ ***R***^*T*×*N*×*C*^ corresponding to *T* frames, *N* muscles, and *C* channels of morphologies, the muscle activity sequence is represented by a tensor **U** ∈ ***R***^*T*×*N*^ corresponding to *T* frames, *N* muscles, and 1 channel of calcium activity. When predicting the larval pose sequence, an initial pose **X**_0_ ∈ ***R***^*N*×*C*^ and a corresponding muscle activity sequence **U**_1:*T*_ ∈ ***R***^*T*×*N*×*C*^ are fed into model to generate the predicted pose sequence 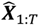; when predicting the larval muscle activity, a final muscle activity state **U**_*T*+1_ ∈ ***R***^*N*^ and a corresponding reversed pose sequence **X**_T:1_ ∈ ***R***^*T*×*N*×*C*^ are fed into model to generate the predicted reversed muscle activity sequence 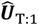

### Software

The code was implemented with Python, the complete project code is available on GitHub: https://github.com/CSDLLab/motor-behavior-recaption. The database that stores the muscle morphological properties (length, width, angle, center coordinates) and muscle activity have been organized into a uniform format to allow for novel analysis.

### Statistics

The Spearmann’s correlation was calculated using Matlab (Mathworks Inc). Other statistics for comparison between groups was performed with prism8.4 and higher version (Graphpad Inc.).

## Acknowledgments

We acknowledge the Bloomington Drosophila stock center for providing the fly stocks, the core facilities of Zhejiang University School of Medicine and the Center of Cryo-Electron Microscopy of Zhejiang University for technical support and reagents. This work was supported by Zhejiang Lab (2020KB0AC02 to Z.G.), the key research and development plan of the ministry of Science and Technology of China (SQ2020YFB130047, 2020YFB1313501), Zhejiang Provincial Natural Science Foundation (LR19F020005), the National Natural Science Foundation of China (31070944, 31271147, 31471063, 31671074, 21874052, 21927802, 61572433 and 61972347) and the Fundamental Research Funds for the Central Universities, China (2017FZA7003).

## Figures

**Figure 1-supplement 1.**
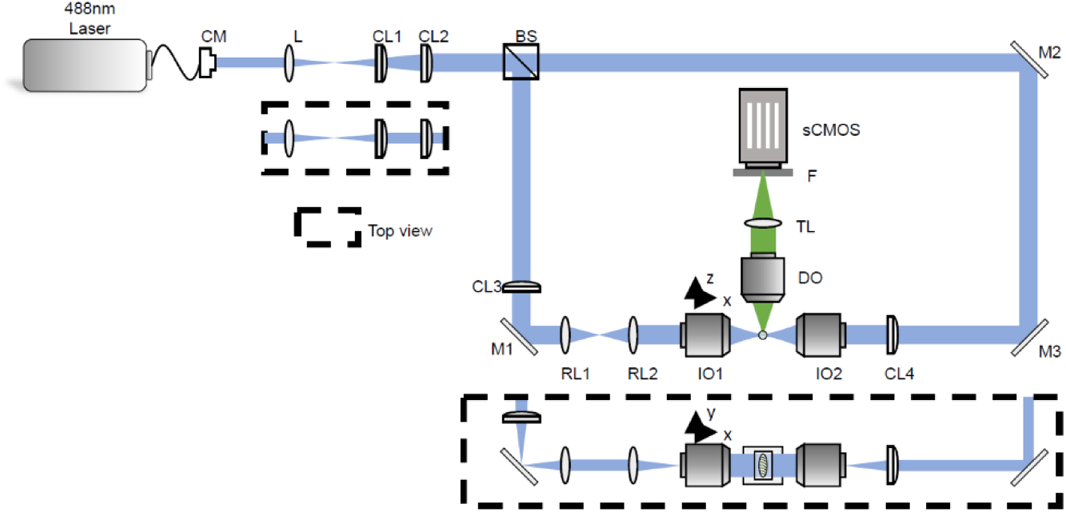
Scheme of the setup of light sheet imaging system. CM, collimator; L, lens; CL, cylindric lens; M, mirror; RL, relay lens; IO, illumination objective; DO, detecting objective; TL, tube lens; F, focal plane.

**Figure 1-supplement 2.**
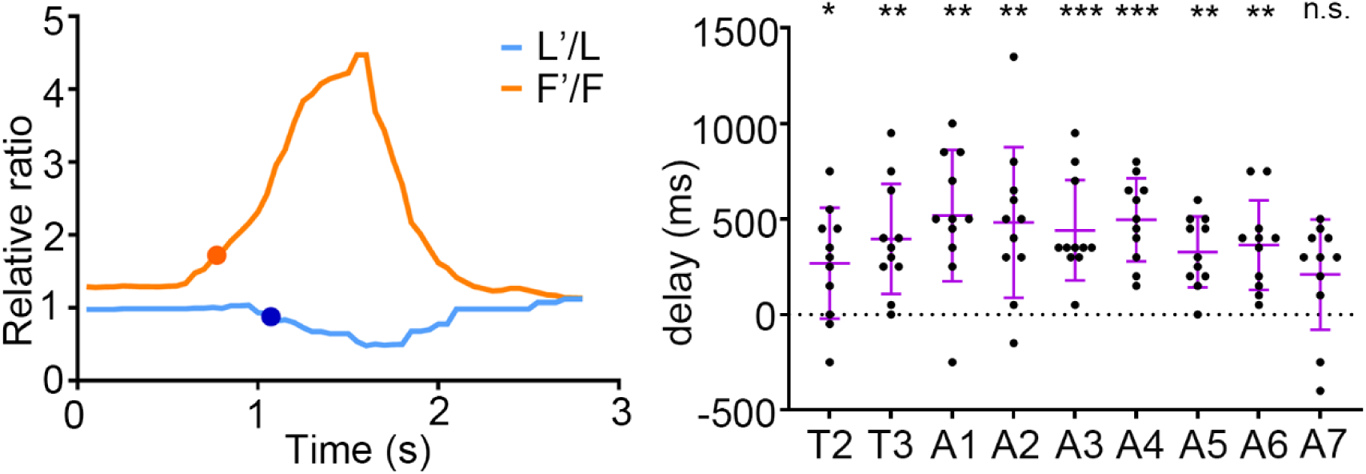
GCAMP marked muscle activity is not caused by muscle deformation. Muscle activation occurs earlier than muscle contraction. A sample curve of muscle activity and muscle length of a larval muscle 9 on left side in segment A1 is shown in left panel. The orange dot and blue dot mark the start time of muscle activation and contraction respectively. Note that activity increase happened earlier than length decrease. Statistics of muscle activity-deformation delay of larval muscle 9 during backward peristalsis is shown in right panel. Muscle activity-deformation delay is always significantly higher than zero except for in segment A7. n=11. n.s. not significant, \**P*<0.05, \*\**P*<0.01, \*\*\**P*<0.001, Wilcoxon Signed Rank test. Error bars, SD.

**Figure 2-supplement 1.**
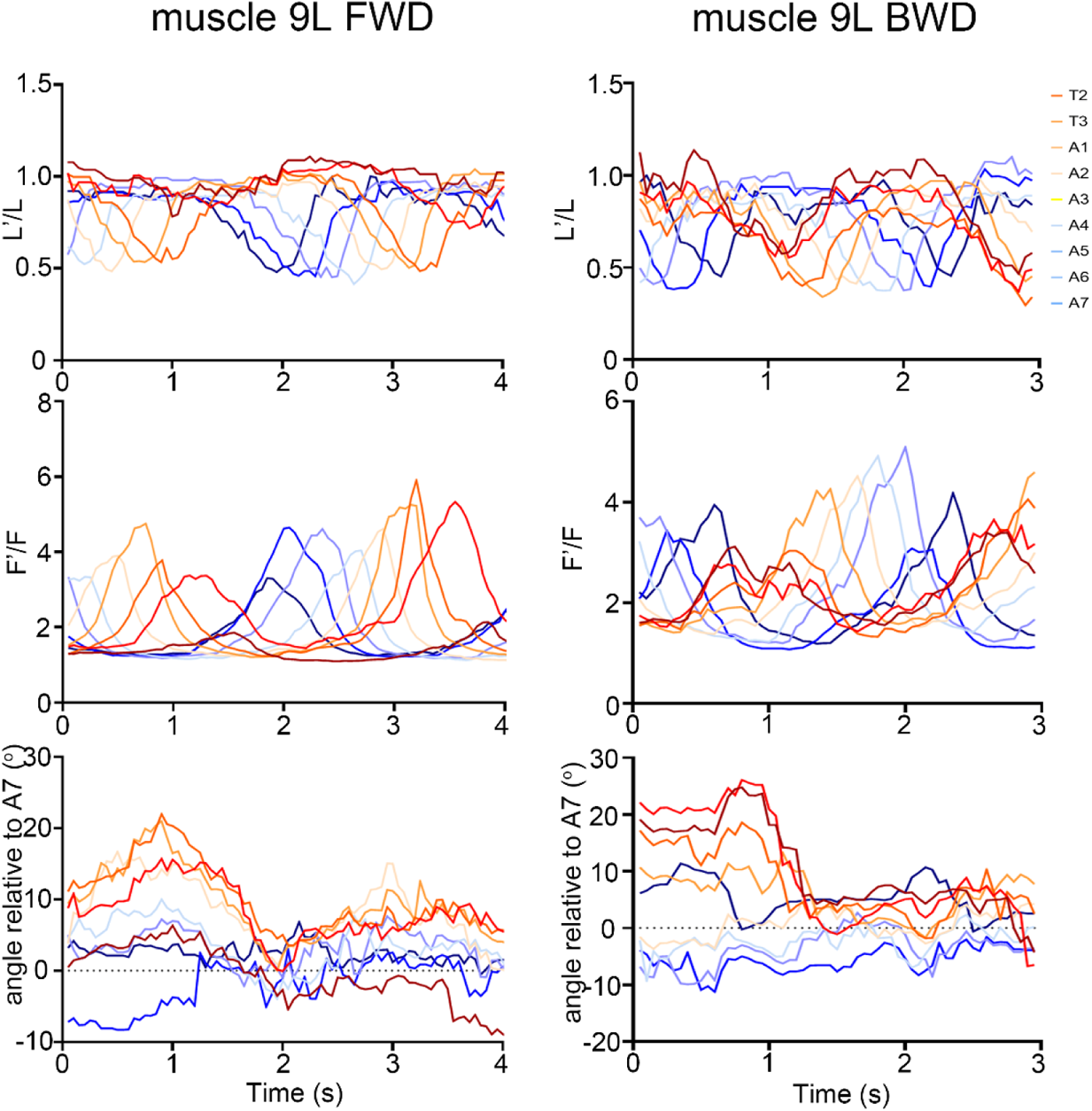
Activity, length and orientation angle of representative dorsal muscle 9 on left during two waves of forward and backward peristalsis.

**Figure 2-supplement 2.**
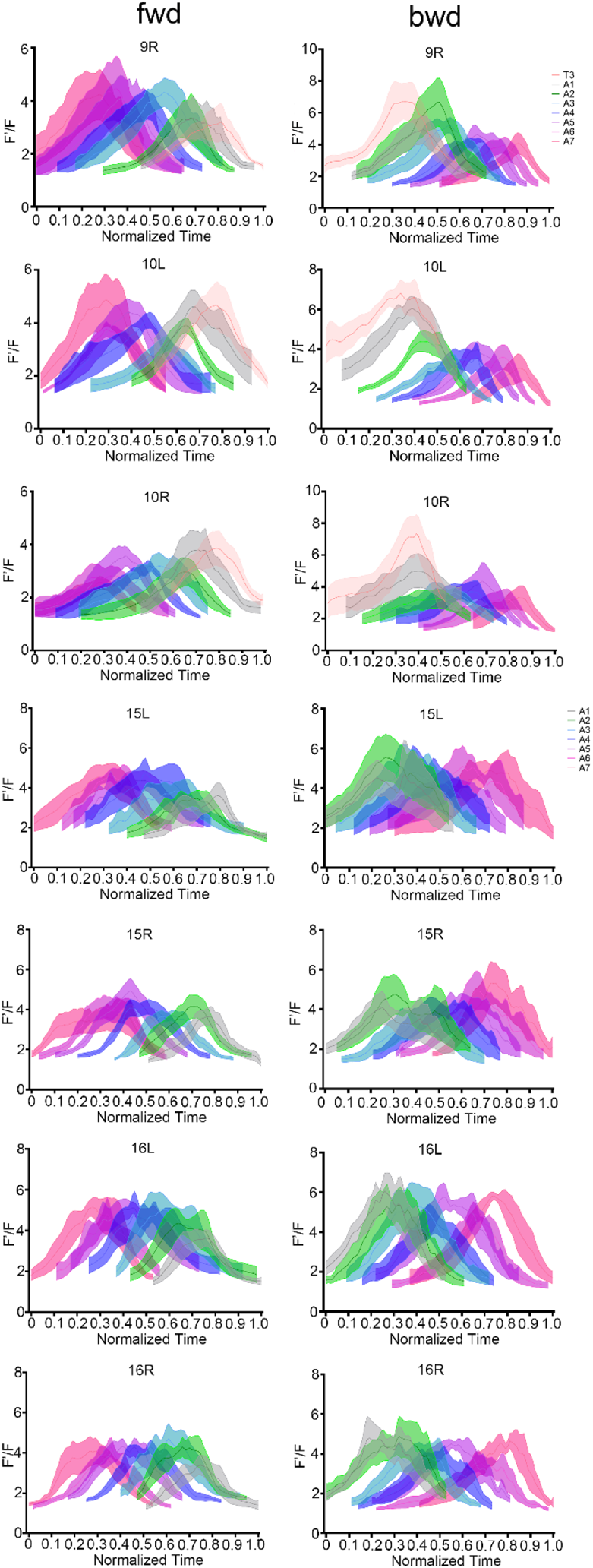
Average normalized one-wave activity pattern of dorsal and ventral muscles in forward and backward peristalsis. For dorsal muscles, n = 6 for forward, n = 5 for backward; for ventral muscles, n = 6 for forward, n = 4 for backward.

**Figure 2-supplement 3.**
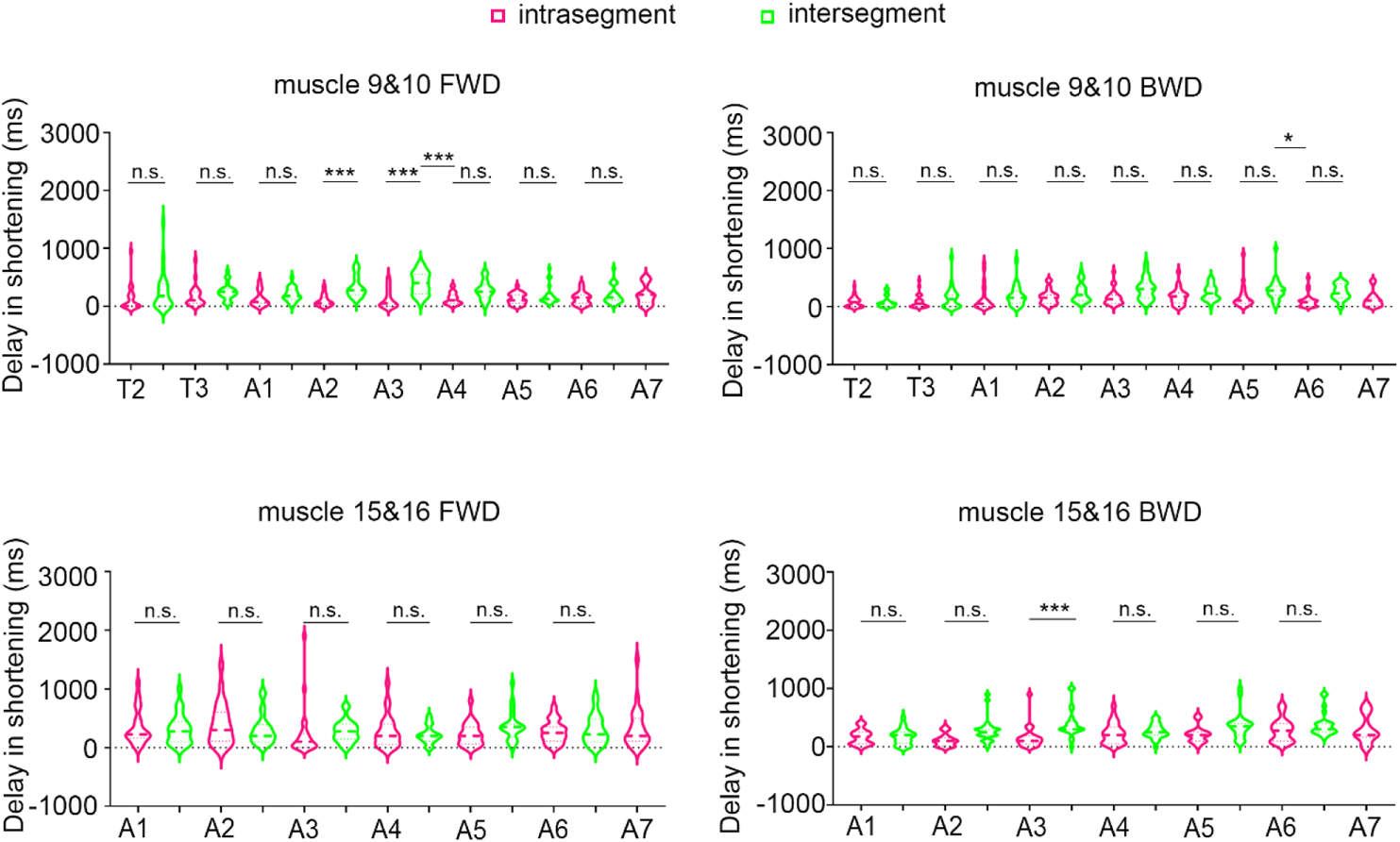
Inter-segmental and intra-segmental delay in shortening of dorsal and ventral muscles in forward and backward peristalsis. The delays within one segment and between neighboring segments are generally not significantly different. Inter-segmental delays and intra-segmental delays are shown in green and red respectively. The labels for intra-segment that is supposed to lie between two neighboring segments are not shown due to limited space. n.s. not significant. * *P*<0.05, ** *P*<0.01, Sidak’s multiple comparison test after two-way ANOVA. n = 24 for all cases. Thick line and dot-line in violin plot, median and interquartile range.

**Figure 2-supplement 4.**
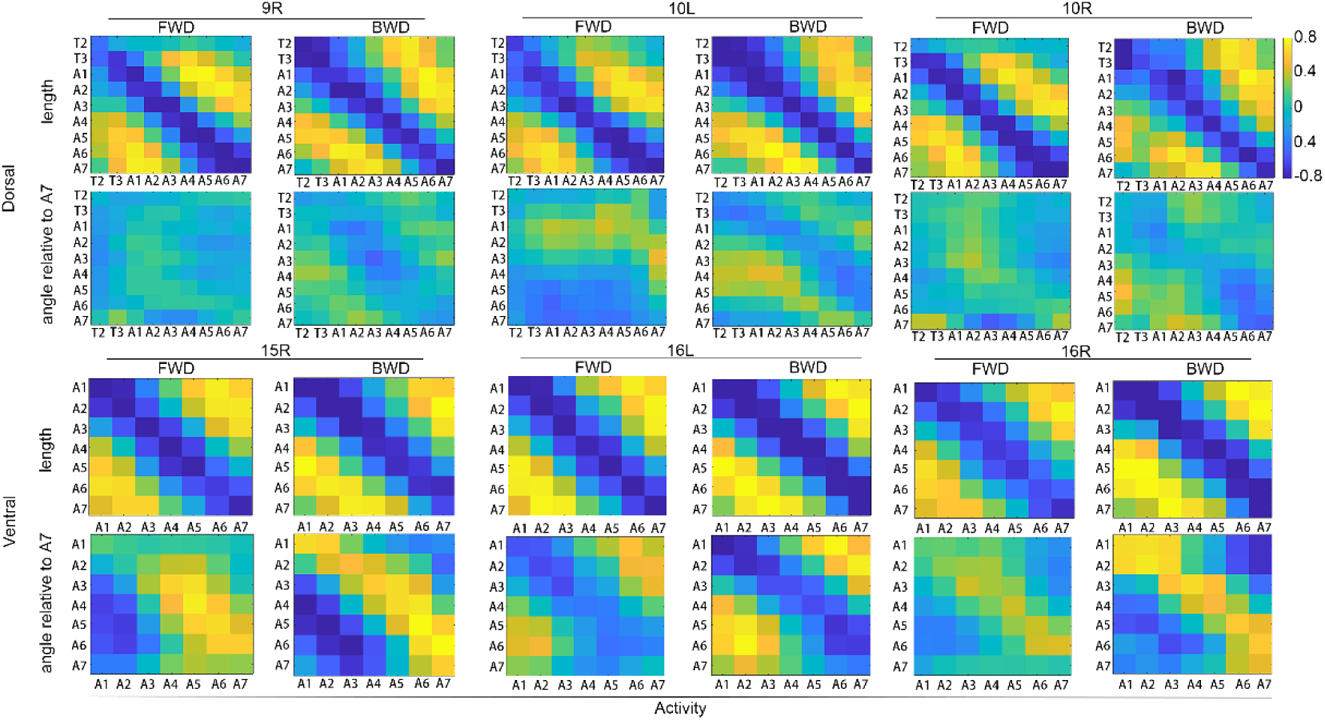
Average correlation between muscle activity and muscle length and orientation angle in dorsal and ventral muscles on left and right in forward and backward peristalsis. A strong negative correlation between length and the activity of the same or neighboring muscle is seen in forward and backward crawling for both dorsal and ventral muscles. Patterns of stronger correlation between activity and orientation angle can be seen in ventral but not in dorsal muscles for both forward and backward peristalsis. Note that the correlation coefficients along the diagonal line in ventral muscles are negative in left muscles and positive in right muscles respectively due to the opposite direction change of ventral muscle orientation angles in left and right during peristalsis. For dorsal muscle 9 and 10, n = 8 waves for forward and n = 6 for backward; for ventral muscle 15 and 16, n = 6 for forward and n = 4 for backward.

**Figure 3-supplement 1.**
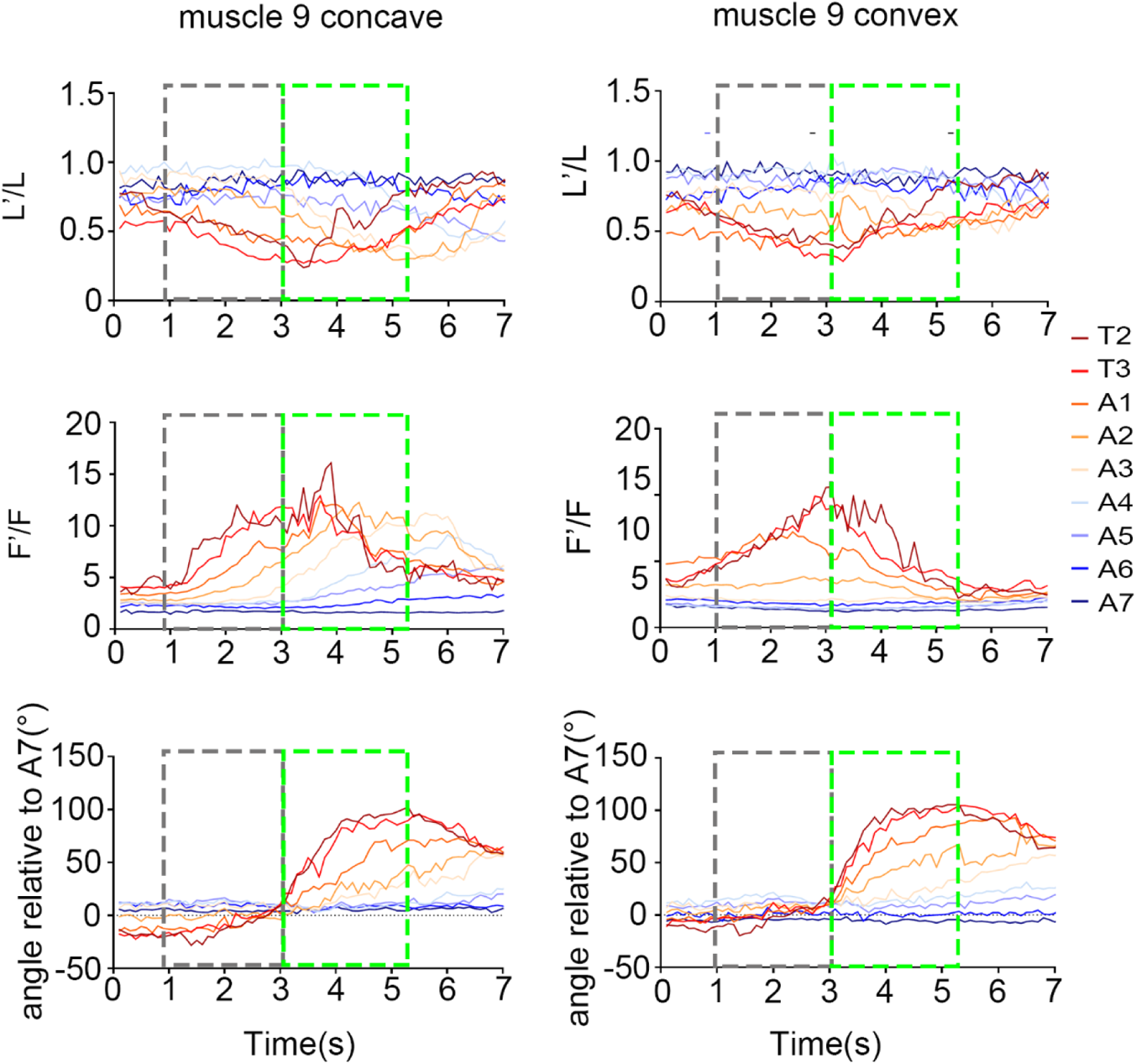
Muscle activity patterns during head cast. Representative curves of activity, length and orientation angle of dorsal muscle 9 on concave and convex side during larval head cast. The grey dotted boxes represent periods of head contraction. The green dotted boxes represent periods of re-orientation in anterior segments.

**Figure 3-supplement 2.**
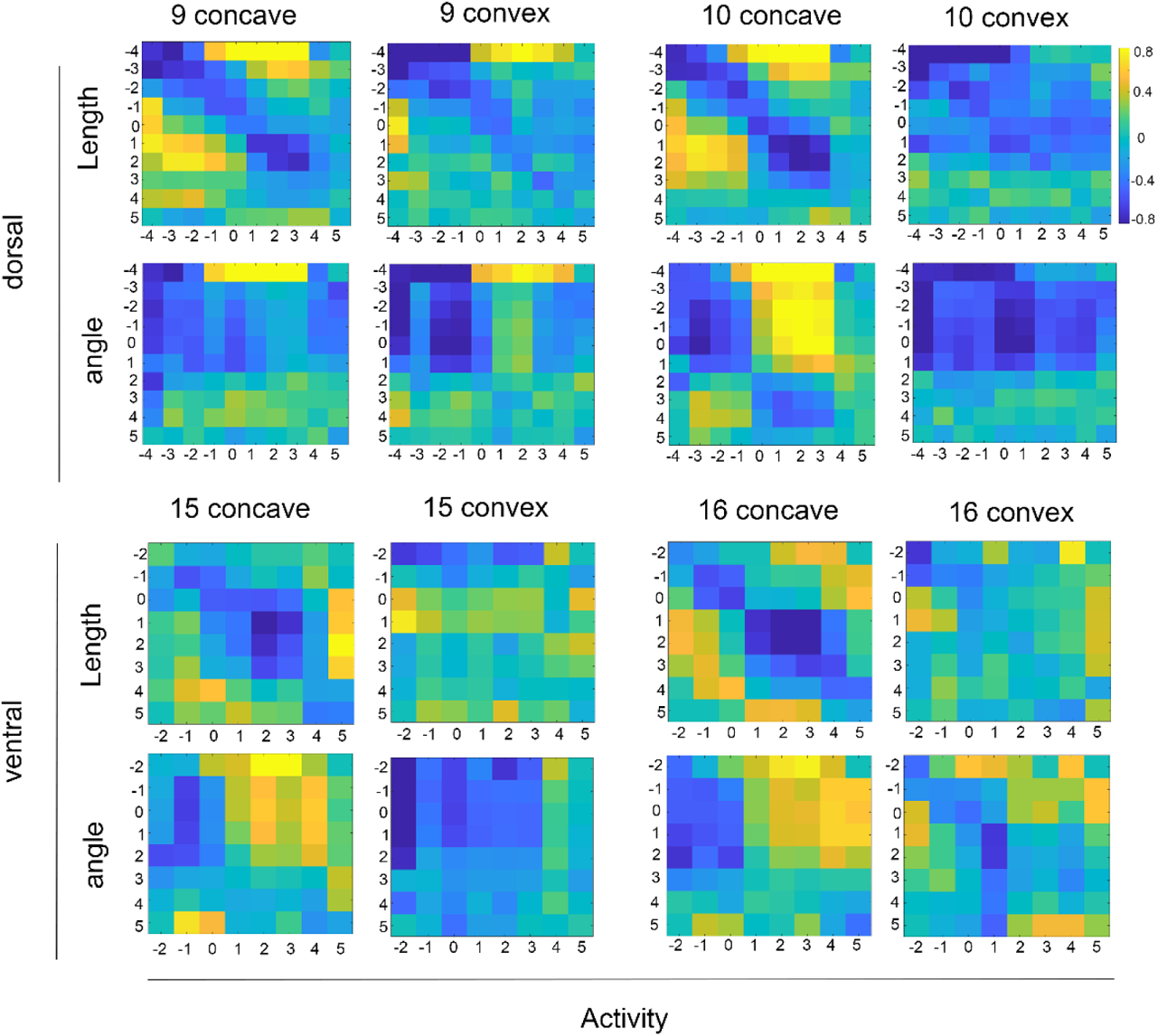
Mean correlation between activity and length and orientation angle of dorsal and ventral on concave and convex side. No consistent map is seen. Segment number 0 is the bending segment. Negative and positive numbers indicate segments anterior and posterior to the bending segment respectively. n=6 for dorsal muscles, n = 5 for ventral muscles.

**Figure 3-supplement 3.**
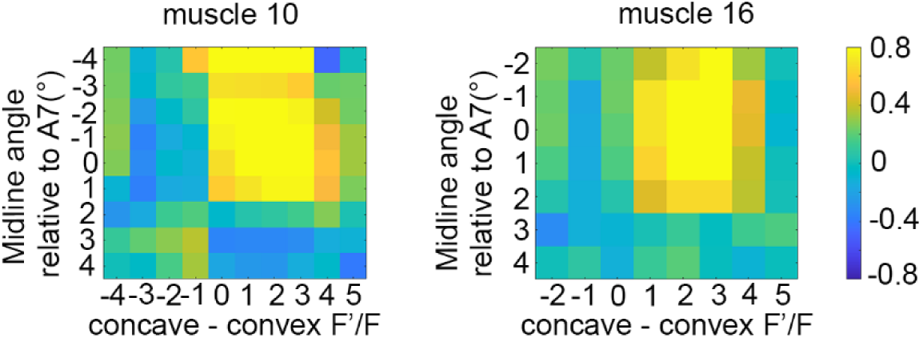
Mean correlation between concave-convex muscle activity difference and segment orientation angle calculated based on muscle 10 (left) and muscle 16 (right). Segment number 0 is the bending segment. Negative and positive numbers indicate segments anterior and posterior to the bending segment respectively. n=6 for muscle 10, n = 5 for muscle 16.

**Figure 3-supplement 4.**
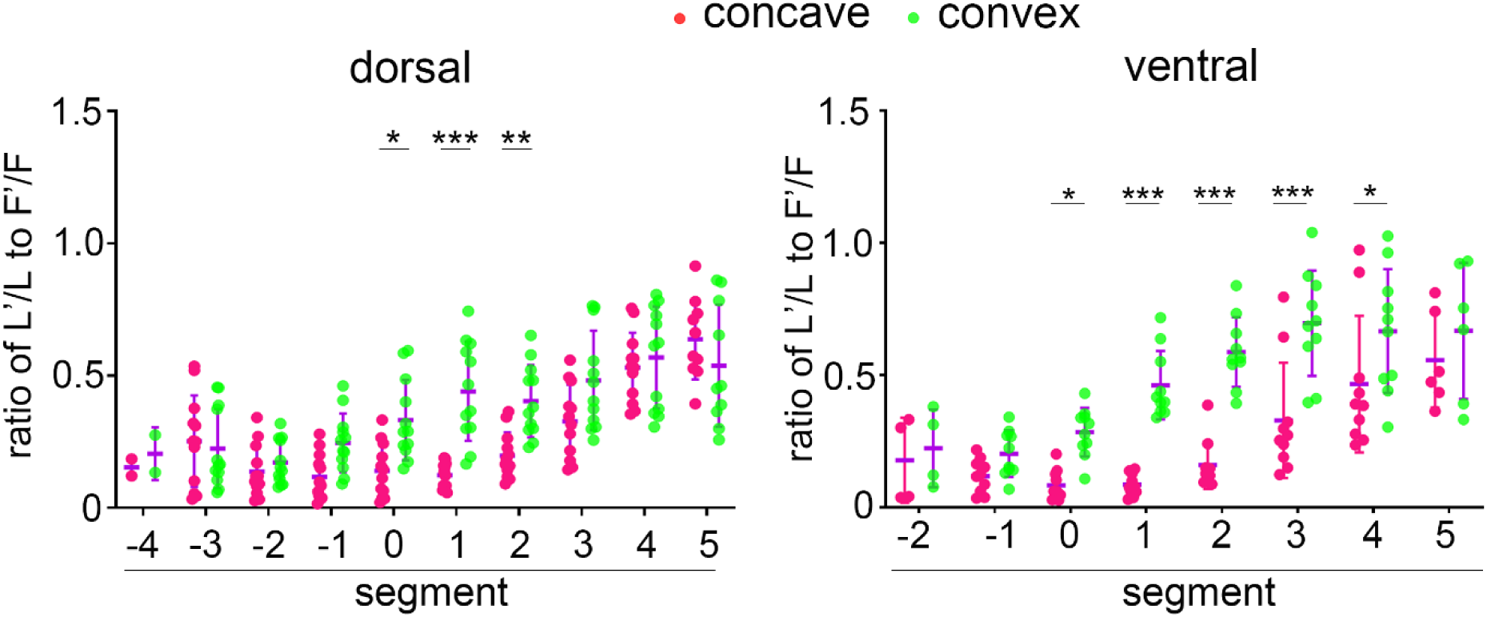
Ratio of muscle length to activity at the time of maximal head cast angle in dorsal and ventral muscles. Segment number 0 indicates the bending segment. Negative and positive numbers indicate segments anterior and posterior to the bending segment respectively. Note that the ratio in convex muscles is generally higher than the concave counterpart in the segment of bending point and more posterior segments in both dorsal and ventral muscles. \**P*<0.05, \*\**P*<0.01, \*\*\**P*<0.001, Sidak’s multiple comparison test after two-way ANOVA. Error bars, SD.

**Figure 4-supplement 1.**
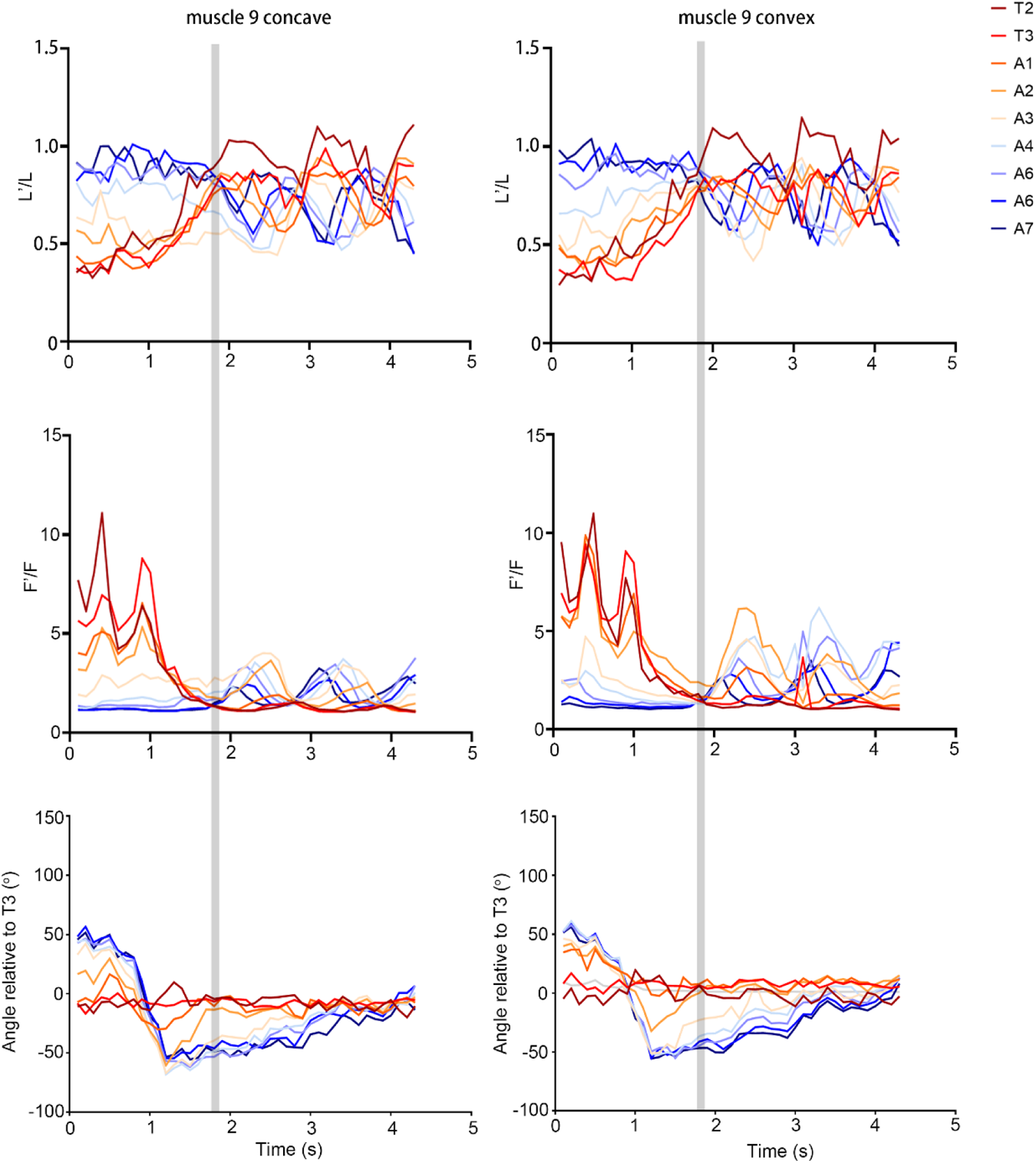
Muscle activity patterns during turning. Representative curves of activity, length and orientation angle of dorsal muscle 9 on concave and convex side during larval turning. The grey lines represent the start of turning after head cast.

**Figure 4-supplement 2.**
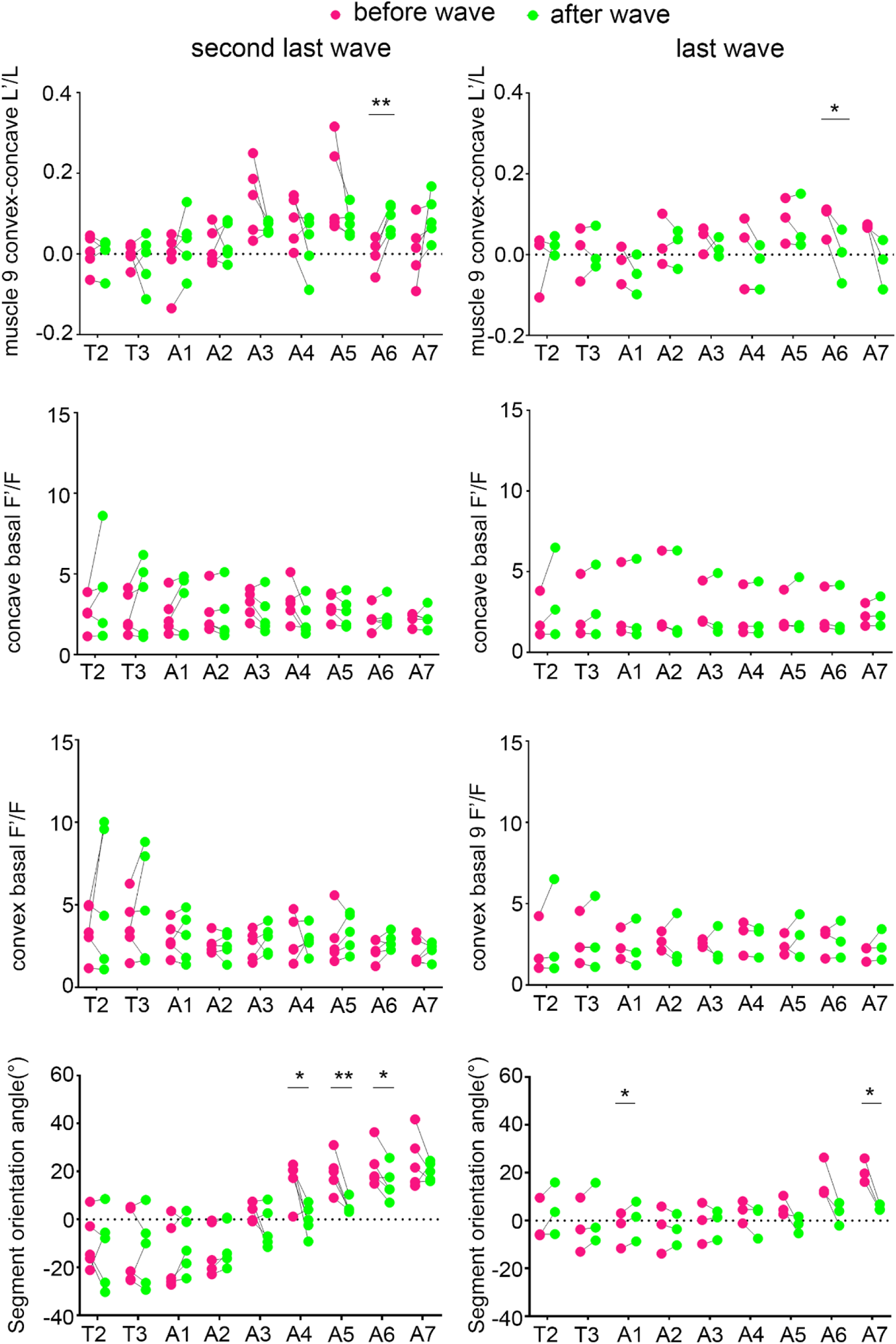
Change of bilateral length difference in dorsal muscle 9, activity of muscle 9 on concave and convex side, and segment midline orientation after the last and second last peristaltic wave during turning. The bending segment is at around A3 before the second last wave and around A5 before the last wave. * P<0.05, ** P < 0. 01, paired *t*-test.

**Figure 4-supplement 3.**
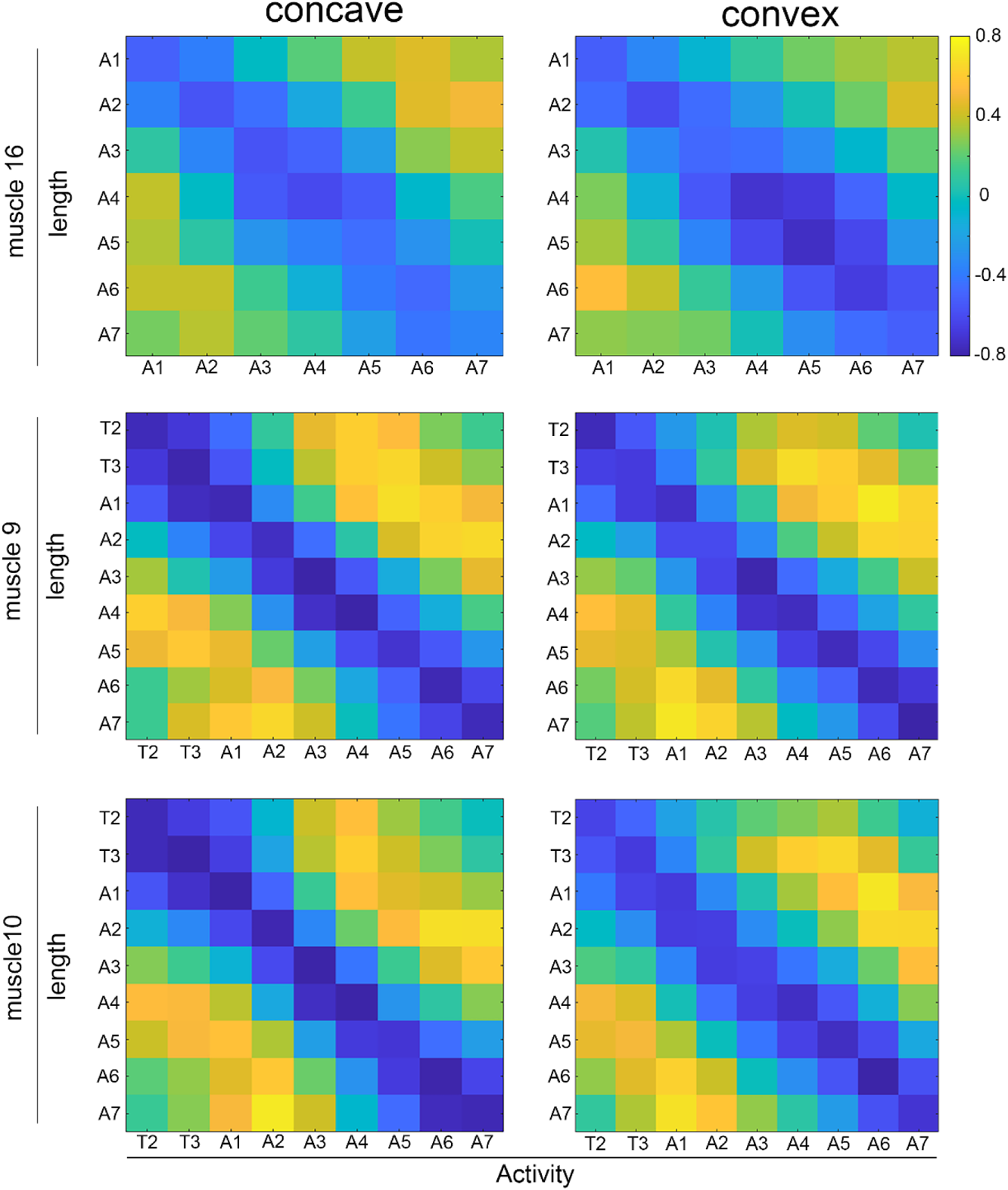
Mean correlation between muscle activity and length of concave and convex ventral muscle 16, dorsal muscle 9 and 10. Strong negative correlation is seen between activity and length of the same muscles. n = 5 for ventral muscle 16. n = 6 for dorsal muscle 9 and 10.

**Figure 4-supplement 4.**
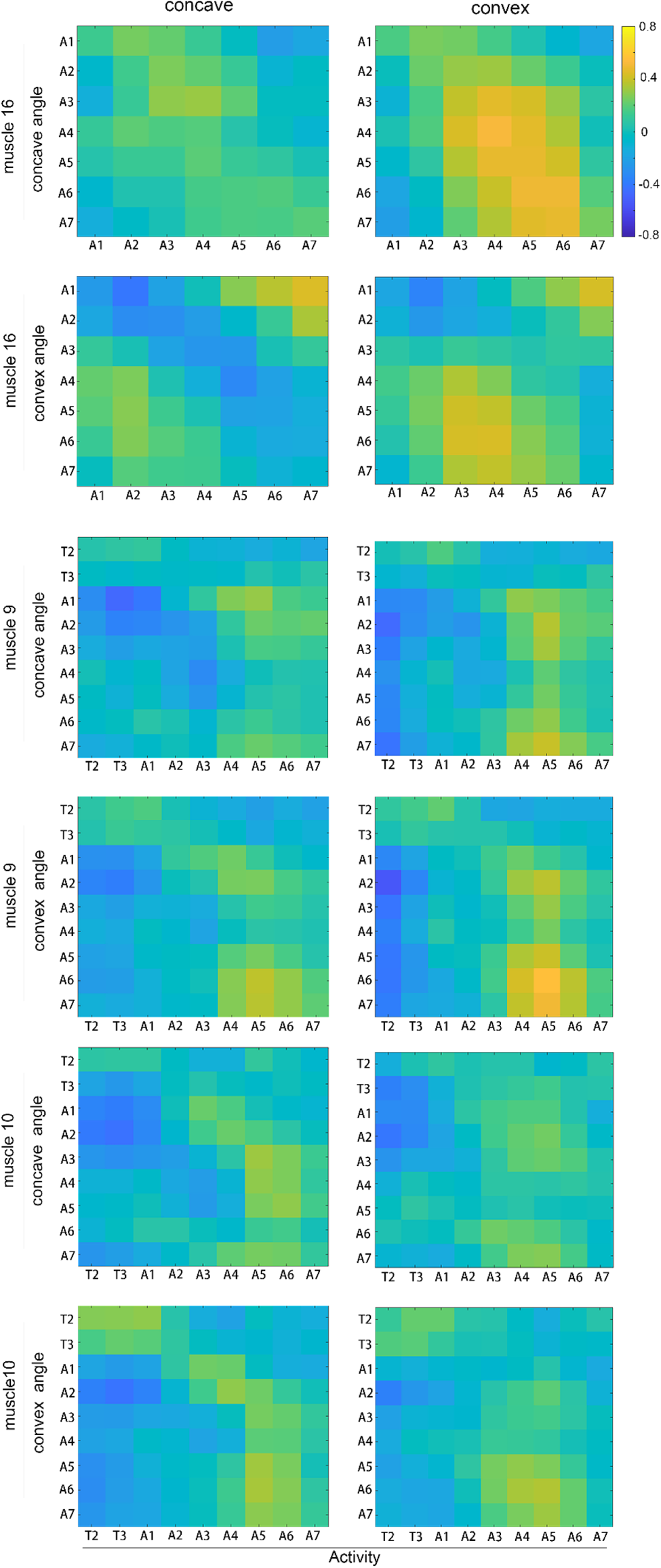
Mean correlation between muscle activity and orientation angle of concave and convex ventral muscle 16, dorsal muscle 9 and 10. n = 5 for ventral muscle 16. N = 6 for dorsal muscle 9 and 10.

## Supplementary File 1A

**Table** Definition of parameters used in the figures.

| Parameters | Description | Figure |
| --- | --- | --- |
| F' | the ratio of the calcium signal of each muscle frame relative to the minimum gray value of the labeled muscle in all labeled frames of the video. | 1f, 1g, 2a, 2b, 2c, 2d, 2f, 3a, 3c, 3d, 3e, 4a, 4d, 4e, 4f, S1b, S2a, S2b, S2d, S3a, S3b, S3c, S3d, S4a, S4b, S4d, S4e |
| $\Delta F' / F'$ | the ratio of the difference between F' and minimal F' to minimal F' | 1f, 1g |
| L' | the ratio of the length of each muscle relative to the maximal length of the muscle 9 left or 16 left in A7 segment in all labeled frames of the video. | 1f, 1g, 2a, 2c, 2d, 2f, 3a, 3e, 4a, 4d, 4e, 4f, S1b, S2a, S2d, S3a, S3d, S4a, S4b, S4c |
| $\Delta L' / L'$ | the ratio of the difference between $L'$ and maximal $L'$ to maximal $L'$ | 1f,1g |
| The original angle | the angle between the long axis of muscle and horizontal axis of the video. | 2a, 2f,<br>3a,3b,3c,3d<br>S2a, S3a, S3c,<br>S2d, S3b |
| The A7 midline angle | the average angle of the bilateral original muscle angle in A7 segment. |  |
| The muscle angle relative to A7 segment | the angle between original angle and A7 midline angle. The positive direction is defined as the right side during peristalsis and the concave during head cast |  |
| Start point of the wave in each segment | the time when the muscle calcium signal rise above the 1.1 times of the minimal value of calcium signals of that muscle | 2b, 2e, 4a, S1b,<br>S2b, S4a |
| End point of the wave in each segment | the time when the muscle calcium signal goes below the 1.1 times of the minimal value of calcium signals of that muscle | 2b,<br>S2b |
| Normalization of | The segments to be normalized consist of A7 to T3 in | 2b, |
| peristalsis | dorsal and A1 to A7 in ventral. The peristaltic wave<br>T2 is not clear and very variable so it is not included.<br>Normalization of the muscle calcium signal curves is done by transforming the curved on the time scale of start point- end point set to be 0 - 1. | S2b |
| The starting points of each muscle length wave | the time when the muscle length goes below 0.9 times of the maximal length of each muscle. | 4b, 4c, 4d, 4e,<br>S1b, S2c, S4b |
| The ending points of each muscle length wave | the time when the calcium signal rise above 0.9 times of the maximal length of each muscle | 4c, 4b, 4d, 4e,<br>S4d |
| The intra-segment delays between start point of muscle activity or muscle length of muscles | 9L-9R; 10L-10R, 9L-10L, 9R-10R. | 2e, S2c |
| The inter-segment delays between start point of muscle activity or muscle length | $9L(n+1) - 9L(n)$ ; $9R(n+1) - 9R(n)$ ; $10L(n+1) - 10L(n)$ ; $10R(n+1) - 10R(n)$. n and n+1 mean the segment and the segment after it. | 2e, S2c |
| Muscle contraction | the length of muscle is less than 0.9 times the maximal | 2c |
|  | length for each muscle |  |
| Muscle tone | the length of muscle is longer than 0.9 times the basal length and the calcium signal is higher than 1.5 times basal calcium signal, that is, the minimal calcium signal for each muscle. |  |
| Muscle relaxed | the length of muscle is more than 0.9 times the basal length and the calcium signal is less than 1.5 times basal calcium signal, that is, the minimal calcium signal for each muscle. |  |
| The ratio of the muscle tone in peristalsis or head cast | dividing the time (frames) of muscle tone with the time (frames) of peristalsis or head cast. | 2d, 3e |
| The start point of head contraction | the time that the concave muscle length less than 0.9 times the maximal length for this muscle in T3 segment | 3a, 3c, S3a |
| The end point of head contraction | the start of head cast. | 3a, 3c, S3a |
| The start of head cast | the time point that T3 segment orientation angle relative to A7 is above 20° in the time window following head contraction. Head cast is considered to occur when the midline orientation angle (relative to | 3a, 3b, 3c, 3d, 3e, S3a, S3b, S3c |
|  | A7 segment) of any head segments (usually T2, T3 and A1) rise from 0° to above 40°. |  |
| The end of head cast | The time point that T3 segment midline orientation angle reaches the maximal level | 3a, 3b, 3c, 3d,<br>3e, S3a, S3b,<br>S3c |
| The bending point of head cast | the posterior most segment with orientation angle relative to A7 above 20° during the head cast. | 3d, 3e, S3b,<br>S3c, S3d |
| The difference between concave and convex | F' in concave muscle – F' in convex counterpart. | 3c, 3d, S3c |
| The start point of turning | the start point of peristalsis when the muscle calcium signal in A7 segment rise to the 1.1 times of the minimal muscle calcium signal level following head cast. | 4a, 4f,<br>S4a, S4c, S4d |
| The end of the turning | the end point of peristalsis when the muscle calcium signal in anterior T3 segment drops below 1.1 times of the minimal muscle calcium signal level following head cast. By the end of turning, the tail is aligned with head. | 4f,<br>S4c, S4d |
| Angle relative to T3 midline | the angle between the muscle or segment and the midline of T3 segment. This parameter is used to | 4a, 4f, S4a,<br>S4d |
|  | measure how much the larval tail is biased from the head before and during turning that aligns tail with head. |  |
| original angle in<br>fig4b,4c | We use original angle but not the angle relative to T3 in larval turning because larva head orientation sometime changes so that original angle change in posterior segments can reflect the movement of larval tail more accurately | 4b, 4c |
| The turning point | the anterior most segment with absolute value of orientation angle (relative to T3) above 20° during turning. | 4c |
| The last wave | the peristaltic wave that makes larval tail completely aligned with head. | 4b, 4e, 4d,<br>S4b |
| The second last wave | the peristaltic wave before the last wave. | 4b, 4e, 4d,<br>S4b |
| Segment orientation<br>angle | To make the orientation angles in each samples look more consistent, the orientation angle change corresponding to tail swing in counterclockwise direction is defined as positive. Add 20 degree to all the orientation angles to make the orientation angle of the head segments to be around 0 degree. | 4d, 4e, S4b |
| Muscle 15 convex-concave L' | The L' before and after the wave are the averages of the respective values within the 5-frame time windows before the start point and after the end point. | 4e, S4b |
| concave basal F' & convex basal F' | The F' before and after the wave are the averages of the respective values within the 5-frame time windows before the start point and after the end point. | 4e, S4b |
| Normalized segment area | the area of each frame calculated by the positions of four corner points relative to maximal area of each segment. | 6c, 6d, 6e |
| Segment contraction rate | the minimal area of each segment / the maximal area of each segment | 6e |
| Number of waves | If the minimal segment contraction ratio of a segment less than 1.1 times of the minimal of all segment contraction of T3 segment, the segment contraction is considered as a peristaltic contraction. The number of segment contraction is the number of peristaltic wave. | 6d |
| The number of head cast | number of head cast after five external stimulations. | 6f |
| The size of head cast | the maximal angle between T3 midline and A7 midline, which must greater than 20 degrees. | 6g |

## Supplementary File 1B

### Supplementary Methods

#### Setup of prediction model

##### Notation and Overview

Larval pose is formed by a set of correlated muscles, which we denote by the graph *G* = {***V***, ***A***}, where ***V*** = {ν_1_, …, ν_*N*_} represents the *N* muscle nodes, ***A*** ∈ *R*^*N*×*N*^ is an adjacency matrix where initially ***A***_*i,j*_ = 1 if an edge directs from ν_*i*_ to ν_*j*_ and 0 otherwise. Each node ν_*i*_ in the graph has a corresponding *C*-dimensional feature vector *x*_*i*_, the entire *T* steps pose sequence feature matrix **X** ∈ *R*^*T*×*N*×*C*^ = {*x*_*t*,*n*_ ∈ *R*^*C*^|*t*, *n* ∈ *Z*, 1 ≤ *t* ≤ *T*, 1 ≤ *n* ≤ *N*}, stacks *T* ×*N* feature vectors. Each channel of node features encodes length *l*, width *w*, angle *a*, and the centroid coordinate (*x*, *y*) respectively for muscle, *i.e.*, the number of node dimensions *C* = 5. Similarly, the entire muscle activity sequence feature matrix **U** ∈ *R*^*T*×*N*×1^ stacks *T* × *N* calcium intensity values.

To accomplish the rational translation between muscle activity **U** and pose sequence **X**, the target sequence generation follows the differential equation:

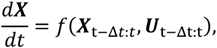

where Δ*t* is a hyper-parameter indicating the length of historical information used in predicting changes at next step. *f* represents a generator neural network that captures the internal interplay patterns between muscles and estimates the effects of muscle activity on pose changes. This generator neural network is implemented with encoder-decoder architecture, and trained in an adversarial manner.

##### Network architecture

There are a generator network and a discriminator network used in training stage. The generator network learns translation between muscle activities and pose sequence, and the discriminator network learns to distinguish whether target sequence is generated by the generator network or sampled from real data. Incorporating the adversarial generator-discriminator training enhance the performance of generator network. The generator neural network consists of encoder and decoder. The encoder network is composed of two layers of spatial-temporal graph convolution operators and one LSTM layer. The decoder is composed of one LSTM layer, two linear layers and two layers of spatial-temporal graph convolution operators. The spatial-temporal graph convolution operator served as low-level feature extraction/reconstruction and the LSTM layer served as long-range nonlinear time series feature modeling. The encoder takes a frame of muscle pose and a set of subsequent muscle activity sequences as input, encodes the joint patterns of muscle activity and initial pose into context features as input of decoder. Conditioned on the context features, the decoder extracts and reconstructs the subsequent pose changes sequence in autoregressive manner. The discriminator network employs the same architecture as the encoder, and is trained to maximize the difference between generated and realistic sequences, whereas the generator network is trained to minimize the difference.

##### Spatial-Temporal graph convolution operator

The spatial-temporal graph convolution stacks two steps: (i) the spatial graph convolution and (ii) the temporal graph convolution. Following Yan al et.^1^ the spatial-temporal graph convolution can be achieved using the node adjacent matrix *A* and entire nodes feature matrix *X* as:

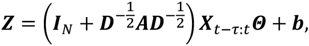

where ***Z*** ∈ ℝ^*N*×*F*^ is the output of spatial neighbors feature aggregation that contains *F* channels features, ***I***_*N*_ ∈ ℝ^*N*×*N*^ denote the identity matrix with *N* dimensions indicating the self-loop of the nodes, ***D*** ∈ ℝ^*N*×*N*^ is the diagonal degree matrix with ***D***_*ii*_ = ∑_*j*_ ***A***_*ij*_, the hyper-parameter *τ* controls the temporal range to be included in the neighbor graph and can also be referred to as the temporal convolution kernel size, *Θ* ∈ ℝ^*τ*×*C*×*F*^ and ***b*** ∈ *R*^*F*^ denote the learnable weights and bias respectively. From the view of one node (e.g., node i at moment *t*), the spatial-temporal graph convolution can be regarded as transforming the features of X_t,i_ ∈ ℝ^1×*C*^ and its spatial-temporal adjacent nodes {**X**_*k,j*_ | ***A***_*i,j*_ ≠ 0, *t* − *τ* ≤ *k* ≤ *t*} to ***Z***_*i*_ ∈ *R*^1×*F*^ with the shared *Θ* and ***b***, in which the shared learnable parameters are useful to learn the most prominent patterns among all nodes in muscles interactions. In practice, a complete spatial-temporal graph convolution operator is implemented with standard 2D convolution and multiplies results with normalized adjacent matrix on the node number dimension.

##### Encoder-Decoder generator neural network

The generator neural network seeks to model the conditional probability of the output sequence given the input initial pose and muscle activity sequence, *i.e.*, *p*(*x*_0_, *u*_0_, …, *u*_*T*−1_).

We applied spatial-temporal graph convolution on each node to capture the correlation of muscles across spatial and temporal dimensions. The features from spatially adjacent nodes and the features from the same node in consecutive poses will be fused and propagated forward repeatedly in a graph convolutional neural network iteratively (illustrated in Figure 6a). After feedforwarding of several layers, the model was capable of capturing the relationship patterns of entire graph sequence.

##### Encoder

The encoder stacks two layers of spatial-temporal graph convolution operators to double the input sequence feature channel for capturing the correlation of muscles across spatial and temporal dimensions. The features from spatially adjacent nodes and the features from the same node in consecutive poses will be fused and propagated forward spatial-temporal graph layers as the input of LSTM layers. Then the LSTM layers were used to model the dynamics of sequential features. Where the original input is combination of a muscle activity sequence feature matrix **U** and an initial pose. We first duplicated the initial pose *T* times to form a sequence, then concatenated two sequences along the feature channel dimension as input. By processing the entire input sequences, the final hidden states of LSTM encoded the overall muscle activity patterns. These hidden states were the initial states of decoder LSTM. The output feature sequence was the input to decoder.

##### Decoder

The decoder stacks one LSTM layer, two linear layers and two layers of spatial-temporal graph convolution operators aims to extract pose dynamics features from the encoder output sequence and reconstruct it into pose sequence. The decoding of decoder works as:

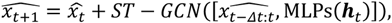

where ***h***_*t*_ is the output of decoder LSTM layer at *t* moment, 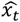 is the generated pose at *t* time step, MLPs(⋅) is stacked two linear layers, [] represents the feature concatenation operation, and *ST* − *GCN*(⋅) is stacked graph convolutional operators for mapping latent features into muscle states displacements (*i.e.*, velocity features) based on historical poses and muscle activity information.

Put it all together, we employed an encoder-decoder architecture to translate a long-term muscle activity sequence into larval pose sequence.

##### Discriminator

To train a model that captures the rich internal dynamics of joint muscle states and external control signals, we used a generative adversarial training approach. The generator neural network is referred to as a *f_G_* and a discriminator *f_D_* for the purpose of simplicity. The discriminator *f_D_* has similar architecture used by the encoder of generator. The ST-GCN is used for high-level features extraction followed by LSTM layers for modeling long term patterns of muscle states and external control signals jointly. Using the discriminator model, the distance between the generated fake data and the real data is as follows:

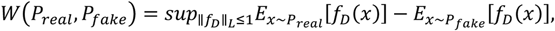

where *P_real_* indicates the distribution of real muscle states, *P_fake_* indicates the distribution of generated muscles states, ‖*f_D_*‖_*L*_ ≤ 1 means that which indicates the discriminator function has to be 1-Lipschitz (or *K*-Lipschitz for some constant *K*).

##### Adversarial Training

Given the generator and discriminator, our LSTM Encoder-Decoder Dynamic System aims to minimize the difference between generated muscle states and realistic muscle states:

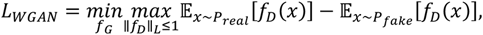

where the adversarial loss *L_WGAN_* equivalent to the Wasserstein distance between the realistic data distribution and the generated fake data distribution, 𝔼[·] indicates the expectation, the generator *f_G_* aims to cheat the discriminator thus minimizing the loss, while the discriminator aims to discriminate the data is real or fake thus maximizing the loss. Besides that, in order to improve the realistic of visual effects of generated muscle states, we added a reconstruction loss for the sequential states reconstruction, applying the L1 loss between all the generated muscle states and realistic muscle states:

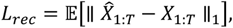

where the 𝔼[·] indicates the expectation, the reconstruction loss *L_rec_* indicates the overall morphological difference between the generated pose sequence 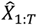 and the realistic pose sequence *X*_1:*T*_.

To this end, the complete training loss is:

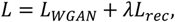

where *λ* weights the reconstruction term.

#### Translation between muscle activities and pose sequences

##### Data preprocess

The muscle states data were collected from individuals Drosophila larvae with calcium imaging, including the muscles on the dorsal side and ventral side. On the dorsal side, 38 clear and bilaterally symmetric muscles were selected. On the ventral side, 30 clear and bilaterally symmetric muscles were selected. The rectangular masks are used in muscle area labeling based on the geometric characteristic of muscles. The dorsal side dataset included 11 individuals and counts over 1200 frames in total. The ventral side dataset included 15 individuals and counts over 1400 frames in total. The datasets were formed by original TIFF image sequences and the corresponding json labels. The muscle area representation was transformed from the original cartesian coordinates of rectangle vertices into length, width, angle, and center coordinates. The average calcium intensity in muscle area was measured as the activity level of the muscle. All calcium activity values were pre-normalized to the range of [0, 1].

##### Implementation

All our models were implemented with Pytorch and trained by WGAN-Clip, using RMSprop optimizer with learning rate 0.0001, the clipping value is 0.01 and the *λ* value is 0.01. To handle the different shapes and calcium levels of individuals, all muscle states and calcium intensities were normalized according to the mean and standard values of the whole dataset.

##### Training

If there are no special instructions, all training is done in generator-discriminator adversarial training manner for 2000 epochs. And the batch size is 4, the lengthes of all training sequences are fixed with *T* = 64 and the sampling gap between consecutive poses is 100ms.

##### Muscle activities to pose sequence translation

Long-term pose sequence prediction aimed to translate muscle activities into corresponding pose sequence over 6400 milliseconds, which was challenging due to accumulated estimation errors and nonlinearity biochemical state variations. Let ***X*** be the morphological features (length, width, angle, and the center coordinates) of pieces of the muscles, and let ***U*** be the muscle activities, and the initial **X**_0_ as arbitrary pose at any moment, then train the model on ventral data and dorsal data separately. For ventral data, the **X** ∈ ℝ^**64×30×5**^ and **U** ∈ ℝ^**64×30×1**^, conditioned on the initial pose **X**_0_ ∈ ℝ^30×5^, the spatial-temporal graph convolution layers of encoder learn to capture the patterns of relationship between individual muscle activities, and the LSTM layer learns to model the long-term muscle activities patterns. Then, in the decoder, the muscle activity patterns from encoder is used to navigate the autoregressive generation of individual muscle hidden features. Finally, with linear layers and spatial-temporal graph convolution layer, hidden features are reconstructed back into morphological features. For dorsal data, the only differences were **X** ∈ ℝ^**64×38×5**^ and **U** ∈ ℝ^**64×38×1**^.

##### Pose sequence to muscle activities translation

In task translating muscle activity sequence into pose sequence, we followed the concept that future pose is determined by past muscle activities. Thus, in task translating pose sequence into muscle activity sequence, we also follow the concept that past muscle activities is determined by future pose. Extrapolating the muscle calcium activity sequence from the pose sequence can be regarded as the rewind of the task of extrapolating the larval pose change sequence from the muscle activity sequence. This is also challenging due to accumulated estimation errors and nonlinearity biochemical state variations. For ventral data, suppose **X** ∈ ℝ^**64×30×1**^ are the muscle activities matrix, **U** ∈ ℝ^**64×30×5**^ are morphological features (length, width, angle, and center coordinates) matrix, and the initial **X**_0_ is muscle activity states at the moment after the last moment of chose muscle activity sequence, then flip the **X** and **U** along the temporal dimension. In the view of training, the only difference between the translation of muscle activity sequence to pose sequence and the translation of pose sequence to muscle activity sequence is the definition of input and output.

##### Pose disparity metrics

To evaluate the appearance difference between estimated muscle behaviors and ground truth muscle behaviors, we transformed muscle representation from node features into cartesian coordinates. Each piece of muscle was converted from morphological features (length, width, angle, center coordinates) to coordinates of four rectangle vertices. We used the Procrustes analysis between the estimated muscle vertices key-points and corresponding ground-truth key-points as appearance comparison metric. In Procrustes analysis, each input matrix was considered as a set of points (the rows of the matrix). The difference between the shape of two poses were evaluated after “superimosing” the two shapes by translating, scaling and optimally rotating them. Then, the square root of the transformed shapes between corresponding points represented the shape disparity,

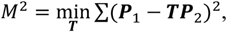

where *M* is the Procrustes analysis disparity, ***T*** is an affine transform matrix, ***P***_1_ and ***P***_2_ are matrices with rows of key points coordinates.

## Notes

### Competing Interest Statement

The authors have declared no competing interest.

